# Tumour architecture inverts ARNT-dependent growth regulation and reveals CPE-mediated spheroid expansion in luminal breast cancer

**DOI:** 10.64898/2026.10.27.761882

**Authors:** Hande Ozkan-Coskun, Ferzan Betül Berber, Elif Kansız, Ezgi Nur Çatı, Mustafa Sami Ozaydın, Zehra Canbulat, Elif Güzar, Begüm Durdar, Dilek Telci, Gozde Korkmaz

**Affiliations:** Research Center for Translational Medicine (KUTTAM), Koç University, Rumelifeneri Yolu, Sarıyer, 34450 Istanbul, Türkiye; School of Medicine, Koç University, Rumelifeneri Yolu, Sarıyer, 34450 Istanbul, Türkiye; Biomedical Sciences and Engineering Program, Koç University, Rumelifeneri Yolu, Sarıyer, 34450 Istanbul, Türkiye; Graduate School of Health Sciences, Koç University, Rumelifeneri Yolu, Sarıyer, 34450 Istanbul, Türkiye; Department of Genetics, Aziz Sancar Institute of Experimental Medicine, Istanbul University, Vakıf Gureba Caddesi, Çapa Campus, Fatih, 34093 Istanbul, Türkiye; Department of Genetics and Bioengineering, Yeditepe University, 26 Ağustos Campus, İnönü Mahallesi, Kayışdağı Caddesi No. 326A, Ataşehir, 34755 Istanbul, Türkiye; Functional Genomics Lab, Koç University, School of Medicine, Istanbul, Türkiye

**Keywords:** Transcriptional regulation, three-dimensional spheroids, tumour architecture, CRISPR screening, ARNT, carboxypeptidase E

## Abstract

Tumour architecture profoundly influences cancer cell behaviour, yet functional genomic studies predominantly rely on two-dimensional (2D) cultures. Here, we performed parallel chromosome 1q-focused CRISPR screens in matched 2D monolayers and three-dimensional (3D) spheroid cultures of luminal breast cancer cells, which revealed ARNT as a prominent architecture-dependent fitness regulator. ARNT depletion reduced proliferation in 2D but enhanced spheroid growth in 3D. This functional switch was reproduced in MCF7 and T47D but not in non-tumorigenic MCF12A cells, indicating a tumour- and architecture-dependent role of ARNT. Transcriptomic profiling revealed distinct responses to ARNT perturbation in 2D and 3D and pointed to carboxypeptidase E (CPE) as a downstream effector. ARNT modulation reciprocally altered CPE expression, and genetic epistasis established CPE as required for ARNT-loss-driven spheroid expansion and sufficient to restore growth in ARNT-overexpressing spheroids. In primary tumours, ARNT and CPE were inversely correlated in low-hypoxia TCGA Luminal A samples, and single-cell RNA sequencing revealed an inverse association within ER-positive cancer cells. Together, these findings demonstrate that the growth-regulatory function of ARNT depends on tumour architecture and that a functional ARNT–CPE axis controls luminal breast cancer spheroid growth, underscoring the importance of incorporating 3D context into functional genomic approaches.

## Introduction

Transcription factors are central regulators of cancer cell survival and proliferation, integrating intracellular signaling and environmental cues into gene expression programs that shape cellular behavior ^1^. However, their regulatory activity is not fixed and may vary according to the cellular environment. Three-dimensional (3D) tissue organization can influence signaling pathways, chromatin organization, and transcriptional activity, potentially altering the molecular requirements for cancer cell growth ^2–5^. Understanding how cellular architecture influences transcription factor function is therefore important for identifying context-dependent mechanisms that support cancer cell fitness.

Functional genomic approaches, particularly CRISPR–Cas9 loss-of-function screening, have enabled the systematic identification of genetic dependencies that sustain cancer cell growth and survival ^6–8^. However, most large-scale functional screens have been performed in two-dimensional (2D) monolayer cultures, which do not fully reproduce the structural and microenvironmental features of tumors ^9,10^. Studies using 3D spheroid models have revealed growth dependencies that differ from those observed in conventional monolayer cultures ^11,12^. More recently, large-scale comparisons of CRISPR screening datasets have further demonstrated that genetic dependencies vary systematically between 2D and 3D growth conditions, particularly among genes involved in cell adhesion, cytoskeletal organization, and metabolic regulation ^13^. These findings indicate that cancer cell fitness is influenced not only by genetic background but also by the architecture in which cells grow.

The effects of cellular architecture are particularly relevant to transcriptional regulators. For example, the mechanosensitive transcriptional coactivators YAP and TAZ respond to changes in cellular organization through altered subcellular localization and transcriptional activity ^14^. Similarly, spheroid-based functional screens have identified NRF2-associated growth dependencies that are not equally apparent under conventional 2D conditions ^12^. Although these studies demonstrate that transcriptional regulation is sensitive to cellular context, it remains unclear whether changes in architecture can reverse the growth-regulatory function of individual transcription factors rather than simply altering their expression or activity. Identifying such regulators and their downstream effectors may provide a better understanding of how cancer cells adapt their growth programs to different cellular environments.

Chromosome 1q is among the most frequently altered genomic regions in breast cancer and contains numerous genes associated with cellular growth and survival ^15,16^. However, the extent to which chromosome 1q-encoded genes contribute to growth dependencies across different culture architectures remains unclear. To address this question, we performed parallel CRISPR–Cas9 loss-of-function screens using a chromosome 1q-focused library in matched 2D monolayer and 3D spheroid cultures of luminal breast cancer cells. This approach identified aryl hydrocarbon receptor nuclear translocator (ARNT; HIF-1β) as a candidate whose functional contribution to cancer cell growth differed markedly between the two culture conditions.

ARNT belongs to the basic helix–loop–helix–PAS family of transcription factors and functions as a central dimerization partner for hypoxia-inducible factor 1α (HIF-1α) and the aryl hydrocarbon receptor (AhR) ^17–19^. Through these interactions, ARNT participates in transcriptional responses to hypoxia and environmental signals, with its regulatory output influenced by partner availability and cellular context ^20,21^. In estrogen receptor-positive (ER+) breast cancer, ARNT has also been implicated in estrogen receptor α (ERα)-mediated transcription and AhR–ERα crosstalk^22,23^. These observations suggest that ARNT can participate in distinct transcriptional programs depending on the signaling environment. Nevertheless, whether cellular architecture alters the growth-regulatory function of ARNT in luminal breast cancer remains poorly understood.

Carboxypeptidase E (CPE) is a multifunctional protein traditionally recognized for its role in the processing and maturation of peptide hormones and neuropeptides^24^. Beyond its established enzymatic activity, CPE has been implicated in cancer cell survival, proliferation, metastasis, and anchorage-independent growth in different experimental models^25–27^. However, the functional contribution of CPE to luminal breast cancer growth, particularly under different culture architectures, remains insufficiently characterized. Whether CPE participates in architecture-dependent transcriptional programs associated with ARNT has not been established.

Here, we identify an architecture-dependent growth-regulatory function of ARNT in luminal breast cancer cells. ARNT loss suppresses proliferation in 2D monolayer culture but enhances spheroid expansion, demonstrating an opposite growth response across culture conditions. Transcriptomic profiling and functional perturbation experiments identify CPE as a downstream effector required for ARNT-loss-associated spheroid expansion, while CPE restoration supports growth in ARNT-overexpressing spheroids. Public chromatin-occupancy datasets indicate ARNT association with the CPE genomic locus, and analyses of bulk and single-cell tumor transcriptomes support an inverse ARNT–CPE expression relationship in selected ER+ breast cancer contexts. Together, these findings establish an architecture-dependent role for ARNT in luminal breast cancer growth and identify CPE as a functional mediator of its three-dimensional growth phenotype.

## Methods

### Cell culture

MCF7 (ATCC HTB-22, RRID:CVCL_0031), T47D (ATCC HTB-133, RRID:CVCL_0553), MCF12A (ATCC CRL-3598, RRID:CVCL_3744), and human embryonic kidney cell line (HEK293T; ATCC CRL-3216, RRID:CVCL_0063) cells were obtained from ATCC and maintained at 37 °C in a humidified atmosphere containing 5% CO₂. MCF7 and HEK293T cells were maintained in DMEM high glucose (Sigma) supplemented with 10% fetal bovine serum (FBS; Biowest) and 1% penicillin– streptomycin (Biowest). T47D cells were cultured in RPMI-1640 (Sigma) with identical supplements. MCF12A cells were cultured in DMEM/F12 (Sigma) supplemented with 5% horse serum (Gibco), 10 ng/ml epidermal growth factor (Gibco), 10 µg/ml bovine insulin (Gibco), 500 ng/ml hydrocortisone (Gibco) and 1% penicillin–streptomycin. All cell lines were tested for mycoplasma contamination at monthly intervals using bioluminescence based MycoAlert® Mycoplasma Detection Kit (Lonza, LT07-318) and only mycoplasma-negative cultures were used in experiments.

### CRISPR library design and pooled library preparation

A custom CRISPR–Cas9 library (LIBGENE1Q) was designed to systematically interrogate genes located on chromosome 1q (Chr1q). The library targets 940 genes, including 80 common essential and 825 non-essential genes, and 33 genes lacking DepMap essentiality classification as defined by DepMap (Public 23Q2; https://depmap.org). Two ribosomal genes (RPL12 and RPS6, located on Chr9) were included as external positive controls, together with 500 non-targeting sgRNAs as negative controls. The complete sgRNA sequence list is provided in Supplementary Table S1.

Pooled oligonucleotide pools encoding sgRNA sequences flanked by cloning adapters were synthesised by Twist Bioscience and PCR-amplified using Phusion® High-Fidelity DNA Polymerase (NEB, M0530L) under the following conditions: initial denaturation at 98 °C for 1 min; 14 cycles of 98 °C for 10 s, 60 °C for 15 s, 72 °C for 30 s; and a final extension at 72 °C for 5 min. Amplicons were purified using the NucleoSpin Gel and PCR Clean-up kit (Macherey-Nagel, 740609) followed by magnetic bead clean-up (CleanNA, CPCR-0050) according to the manufacturers’ instructions.

The lentiCRISPR v2 vector (Addgene #52961; a gift from Feng Zhang) was linearised with BsmBI-v2 (NEB, R0739) by incubating 5 µg of plasmid with 5 µl of BsmBI-v2 at 55 °C for 3 h. sgRNA amplicons were assembled into the linearised lentiCRISPR v2 backbone using the Gibson Assembly® Master Mix (NEB, E2611L). Ligation products were electroporated into electrocompetent Endura cells (Lucigen, 60242) using a MicroPulser electroporator (Bio-Rad). Multiple parallel transformations were pooled to maintain library complexity (>500-fold representation per sgRNA at the bacterial stage). Plasmid DNA was extracted using the M/N Midi-Prep Purification Kit (Macherey-Nagel, 740410).

Library distribution was assessed by amplicon sequencing of the plasmid pool. Sequencing reads were processed using MAGeCK count (version 0.5.9.5), with each sgRNA represented at >100-fold coverage at this stage, confirming uniform sgRNA representation and library complexity.

### Lentivirus production and transduction

LIBGENE1Q library (or 10 µg of individual plasmid for validation experiments), 3.5 µg of pVSV-G (Addgene #8454), 5 µg of pMDLg/pRRE (Addgene #12251) and 2.5 µg of pRSV-Rev (Addgene #12253) was prepared in serum-free DMEM. Transfection was performed using polyethyleneimine (PEI; Sigma, 408727) at a 3:1 (w/w) PEI-to-DNA ratio. The transfection mix was incubated for 15 min at room temperature and added dropwise to cells. After 16 h, the medium was replaced with fresh DMEM containing 10% FBS, and viral supernatants were harvested at 48 h post-medium change, filtered through a 0.45 µm Steriflip-HV PVDF filter (Millipore) and stored at −80 °C until use.

For lentiviral production, HEK293T cells were seeded at 4 × 10⁶ cells per 100-mm dish one day prior to transfection. Per dish, a plasmid mixture comprising 10 µg of the Functional viral titres were determined for each batch by limiting-dilution transduction of target cells followed by puromycin selection, to achieve a multiplicity of infection (MOI) < 0.4, ensuring predominantly single-guide integration per cell. For all transductions, target cells were incubated with viral supernatants in the presence of 8 µg/ml polybrene (Sigma, TR-1003) for 24 h, after which the medium was replaced. Transduced cells were selected with puromycin at 2 µg/ml (MCF7), 4 µg/ml (T47D) or 1.2 µg/ml (MCF12A) for 48 h, or with blasticidin at 5 µg/ml where indicated.

### Pooled CRISPR screening in 2D and 3D culture

Pooled CRISPR loss-of-function screens were performed in MCF7 and T47D cells under matched 2D monolayer and 3D spheroid conditions. Cells were transduced with the LIBGENE1Q library at MOI < 0.4 and a coverage exceeding 1,000-fold per sgRNA. After 24 h, the medium was replaced and cells were selected with puromycin (2 µg/ml MCF7; 4 µg/ml T47D) for 48 h. Following selection, cells were maintained for an additional 48 h to allow elimination of guides causing acute cytotoxicity. A baseline reference fraction (T = 0) was collected at this point, and the remaining cells were split into parallel 2D and 3D cultures and propagated for an additional 20 days, maintaining a minimum of 1,000-fold library coverage at each passage. Endpoint samples (T = 20) were collected for both arms. Three biological replicates were performed per cell line per condition. All cell pellets were stored at −80 °C until DNA extraction. Frozen cell pellets were first thawed on ice, and genomic DNA (gDNA) was extracted using the Macherey-Nagel™ NucleoSpin™ Tissue Isolation kit (Macherey-Nagel, 740952) according to the manufacturer’s instructions.

sgRNA sequences were amplified from gDNA in a two-step PCR strategy. In the first PCR, sgRNA cassettes from integrated lentiCRISPR-LIBGENE1Q constructs were amplified using primer pairs flanking the guide region labelled as PCR1 (Supplementary Table S2). For each sample, sufficient gDNA was used to maintain ≥500-fold sgRNA coverage, with a maximum of 1 µg of gDNA per 25 µl reaction and up to 40 µg of gDNA per sample distributed across multiple parallel reactions; products were pooled prior to second-round amplification. In the second PCR, Illumina P5/P7 adapters and sample-specific indices were appended using 5 µl of first-PCR product per 25 µl reaction (PCR2; primers in Supplementary Table S2). Phusion High-Fidelity DNA Polymerase (NEB, M0530L) was used in PCR and were limited to 15 cycles to minimise amplification bias. Final amplicons were purified by magnetic bead clean-up (CleanNA), pooled at equimolar ratios, and sequenced on Illumina NovaSeq 6000 platform.

### Library quality control and mapping analysis

Library content and mapping ratios were assessed from sgRNA count tables and count-summary files generated by MAGeCK ^28^ (version 0.5.9.5) for the plasmid library and cell line–specific libraries. All analyses and visualizations were performed in Python 3.12 using pandas 2.2.2, numpy 2.0.2, matplotlib 3.10.0 and seaborn 0.13.2. For mapping-quality analysis, count-summary tables were used to derive mapped and unmapped read fractions for each sample. Read-count and mapping-percentage distributions were further summarized using histograms. To evaluate sgRNA representation across the full library, raw count tables were grouped by annotated target gene, genes were ordered according to their mean abundance across samples, and the data were reshaped into long format. Read counts were transformed as log10(count + 1), and whole-library representation was visualized as scatter plots showing gene order against log-transformed read count, with points colored by sample identity.

### CRISPR screen data analysis

CRISPR screen sequencing data were analyzed using MAGeCK ^28^ (version 0.5.9.5). sgRNA-level read counts were generated from FASTQ files using MAGeCK count, with the sgRNA library file supplied via the -l option to define guide identifiers, sequences and target-gene annotations for plasmid and input library.

Separate MAGeCK count tables were generated for each cell line. For MCF7, samples comprised t0, 2D t10, 2D t20 and 3D t20 conditions across three biological replicates; a 3D t10 condition was not included for this line. For T47D, all five conditions were collected: t0, 2D t10, 3D t10, 2D t20 and 3D t20 across three biological replicates.

Differential analyses were performed with MAGeCK test using raw count files. For each comparison, treatment and control groups were specified explicitly using MAGeCK -t and -c arguments. For time-course comparisons, t0 replicates were used as controls and later time-point replicates were used as treatments. For culture-condition comparisons, 2D and 3D groups were compared within the corresponding cell line and matched time points where available.

Pairwise comparisons were performed between all available time points and culture conditions within each cell line. For T47D, this included comparisons against t0, matched 2D versus 3D comparisons at each time point, and longitudinal comparisons within each culture condition. The same comparisons were performed for MCF7 where data were available; comparisons involving 3D t10 were omitted for this line.

Normalization was performed during MAGeCK testing using both non-targeting control sgRNAs and copy-number correction. Non-targeting sgRNAs were supplied as a control-guide list using the --control-sgrna option, together with --norm-method control, so that normalization factors were estimated from the control sgRNA set. Copy-number-aware analyses were also performed using --cnv-norm with the CCLE gene-level copy-number file CCLE_copynumber_byGene_2013-12-03.txt and the corresponding MAGeCK --cell-line argument.

For each cell line, dropout candidates were defined by comparing T = 20 versus T = 0 sgRNA abundances. Candidate depletion events were selected using thresholds of FDR < 0.1 and median sgRNA log_2_fold-change < −0.8. To identify architecture-dependent enrichment, T = 20 sgRNA abundances in 3D culture were compared with the matched 2D condition using MAGeCK, corresponding to a 3D/2D ratio. Enriched candidates were selected using thresholds of FDR < 0.25 and log_2_fold-change > 0.75. These candidate-level comparisons were evaluated at the gene level using the median sgRNA score, with three biological replicates per arm.

To generate density plots, sgRNA count distributions were analysed from count tables generated by MAGeCK using a custom workflow written in Python 3.12 with pandas 2.2.2, NumPy 2.0.2, seaborn 0.13.2 and matplotlib 3.10.0. Sample names were parsed programmatically according to dataset-specific conventions to recover experimental metadata, including time point, culture condition and replicate number. Where needed, parsing rules were adapted to distinguish experimental time labels from other numeric identifiers embedded in sample names. Samples sharing the same experimental attributes were grouped together, and sgRNA abundances were transformed as log_2_(count + 1) before visualization. Distributions were represented as replicate-level kernel density estimates, with replicates plotted in a fixed numerical order and displayed using consistent formatting across panels to preserve comparability.

To determine essential shifting, cumulative density plots were generated from MAGeCK sgRNA summary tables using a custom analysis workflow written in Python 3.12 with pandas 2.2.2, NumPy 2.0.2, Matplotlib 3.10.0 and seaborn 0.13.2. Each MAGeCK sgRNA summary file corresponding to an individual screening comparison was processed separately. sgRNA-level log_2_fold-change (LFC) values were aggregated at the gene level by calculating the mean LFC for all sgRNAs assigned to each gene. Gene-level fold-change values were subsequently integrated with an external essentiality reference table derived from DepMap (Public 23Q2; https://depmap.org), which was used to assign genes to common essential and nonessential categories. Non-targeting guides were treated as a distinct internal control group and were reassigned to a dedicated NT category. Genes that could not be matched to an essentiality class were excluded from visualization. Within each category, genes were sorted by increasing average log_2_fold-change and used to construct cumulative density distributions, enabling comparison of the global distribution of screening effects across reference gene classes and non-targeting controls. A fixed plotting order and color scheme were maintained across all comparisons to ensure visual consistency.

Waterfall plots were generated from MAGeCK output tables using a custom analysis workflow written in Python 3.12 with NumPy 2.0.2, pandas 2.2.2, matplotlib 3.10.0, seaborn 0.13.2, openpyxl 3.1.5 and scipy 1.16.3. For each comparison, gene-level results were parsed from MAGeCK gene_summary files, and gene identifiers together with the corresponding log_2_fold-change values were reformatted into simplified Excel tables for plotting. Genes were then sorted by increasing log_2_fold-change and assigned a plotting rank based on this order, rather than retaining the original MAGeCK rank columns. The resulting rank order defined gene positions in the waterfall plots, with the most depleted genes appearing at the beginning of the distribution. Additional annotation fields were included when needed to enable highlighting and visualization-specific grouping of selected genes.

### Individual sgRNA cloning and validation constructs

For functional validation, individual sgRNAs targeting candidate genes were cloned into lentiCRISPR v2 (Addgene #52961) or lentiCRISPR v2-Blast (Addgene #83480, gift from Mohan Babu) using BsmBI-v2 digestion and ligation of annealed sense/antisense oligonucleotides (T4 DNA Ligase, NEB, M0202). Oligonucleotide sequences are listed in Supplementary Table S3 (Macrogen, USA). Constructs were verified by Sanger sequencing (Macrogen) prior to use.

For ARNT overexpression (ARNT OE), pLX304-ARNT (Addgene #203578; a gift from Abhishek Chakraborty and William Kaelin) was used. The corresponding empty-vector control (pLX EV) was generated by digesting pLX304-ARNT with NdeI and SalI (NEB), blunting with T4 DNA polymerase (NEB, M0203) and self-ligating with T4 DNA ligase.

For functional rescue experiments, a CRISPR-resistant ARNT cDNA (PAM-mutant ARNT) was generated by site-directed mutagenesis of pLX304-ARNT using the Q5® Site-Directed Mutagenesis Kit (NEB, E0554) with the primers (ARNT_SDM_PAMMUT_FWD and ARNT_SDM_PAMMUT_REV) listed in Supplementary Table S4. The protospacer-adjacent motif (PAM) sequence adjacent to the ARNT sgRNA #3 target site was altered from CGG to CGC, abolishing CRISPR– Cas9 cleavage of the construct without changing the encoded amino acid sequence. The mutated construct was verified by whole plasmid sequencing (Macrogen).

For CPE overexpression (CPE OE), the full-length CPE coding sequence (NM_001873) was amplified from MCF7 cDNA using primers carrying PacI and NotI restriction sites as CPE_OE_PRIMER_FWD and CPE_OE_PRIMER_REV (Supplementary Table S4) and cloned into pLJC2-GPT2-3xFLAG (Addgene #163447; a gift from Jason Cantor) digested with the same enzymes, using T4 DNA Ligase. The corresponding pLJC2 empty vector was generated by digesting pLJC2-3xFLAG with NheI and NotI (NEB), blunting with T4 DNA polymerase and self-ligating. Lentiviruses for individual constructs were produced as described above.

### Generation of knockout and overexpression cell lines

For stable knockout (KO) cell line generation, MCF7, T47D and MCF12A cells were transduced with lentiCRISPR v2 (puromycin selection) or lentiCRISPR v2-Blast (blasticidin selection) carrying a non-targeting control sgRNA (CNT) or a target-specific sgRNA (sgARNT, sgCPE, sgLXN or sgARNT INT). Following antibiotic selection, populations were used as polyclonal pools rather than single-cell clones unless stated otherwise. Knockout efficiency was verified by RT-qPCR and immunoblot at the population level prior to use in functional assays.

For double-knockout (DKO) experiments, cells were sequentially transduced — first with lentiCRISPR v2-Blast (blasticidin-selectable) carrying sgARNT or non-targeting CNT-sgRNA, then with lentiCRISPR v2 (puromycin-selectable) carrying sgCPE or non-targeting CNT sgRNA — yielding four genotype combinations: CNT/CNT, sgARNT/CNT, CNT/sgCPE and sgARNT/sgCPE.

For stable overexpression (OE) lines, cells were transduced with pLX304-ARNT, pLJC2-CPE, or the corresponding empty-vector controls (pLX EV, pLJC2 EV) and selected with blasticidin (5 µg/ml) or puromycin (2 µg/ml), as appropriate. For combined OE experiments, cells were sequentially transduced with both ARNT and CPE overexpression constructs and selected with the corresponding antibiotics. Overexpression status was verified by RT-qPCR.

### Intergenic targeting control

To distinguish ARNT gene-specific phenotypes from regional copy-number effects, an intergenic sgRNA (ARNT-INT) was designed to target a non-coding region within ∼10 kb of the ARNT locus, lacking annotated regulatory elements (no overlap with H3K27ac, H3K4me3 or DNase hypersensitivity peaks in ENCODE breast tissue datasets). The ARNT-INT sequence is provided in Supplementary Table S3. MCF7 cells were transduced with lentiCRISPR v2 carrying sgARNT, ARNT-INT, or non-targeting sgRNA, and assessed for ARNT expression (by RT-qPCR) and 3D spheroid growth in parallel.

### GFP competition assay

GFP-expressing reference populations of MCF7, T47D and MCF12A cells were generated by transduction with pLenti-CMV-GFP-Puro (Addgene #17448; a gift from Eric Campeau and Paul Kaufman), followed by puromycin selection. For competitive proliferation assays, GFP-positive reference cells were mixed with sgRNA-transduced (non-GFP) cells at a 30:70 ratio (GFP:sgRNA) and co-cultured under 2D conditions ^29^. The proportion of GFP-positive cells was measured every 72 h for 12 days using a flow cytometry, acquiring a minimum of 10,000 events per sample. Three biological replicates were performed per condition. Relative fitness was calculated as the GFP-positive fraction at each time point divided by the GFP-positive fraction at T = 0.

### Colony formation assay

Cells stably expressing gene-targeting or non-targeting sgRNAs (or OE/EV constructs) were seeded at 4,000 cells per well in 6-well plates in technical triplicate, with three independent biological replicates per condition. Cells were cultured under standard conditions for 10–14 days, with medium replacement every 3 days. At endpoint, cells were washed twice with DPBS, fixed with 100% ice-cold methanol for 20 min at 4 °C, and stained with 0.5% (w/v) crystal violet (Sigma) in 25% methanol for 1 h at room temperature with gentle agitation. Excess dye was removed by extensive washing with distilled water; plates were air-dried and scanned. Total colony area was quantified using ImageJ (Fiji) by applying the same intensity threshold across all wells of a plate and normalised to the average of the corresponding control wells (set to 100%).

### 3D spheroid growth assay and quantification of spheroid-forming-ability

Three-dimensional (3D) spheroid cultures were established by seeding 7.5 × 10⁵ cells onto UV-sterilised plates in final growth medium supplemented with 10% (w/v) methylcellulose. Spheroid formation was monitored over time, and brightfield images were acquired on day 4 and day 6 using a Nikon Eclipse Ts2 inverted microscope with a 20×/0.40 objective. For each biological replicate, at least 50 spheroids were imaged, and three independent biological replicates were performed per condition.

Spheroid size was quantified using ImageJ (Fiji). The spatial scale was calibrated using the microscope scale bar with the *Set Scale* function, and individual spheroids were segmented using the *Polygon Selection* tool. Spheroid area was calculated for each segmented object and reported as fold change relative to the matched control condition.

To assess spheroid-forming ability, equal volumes of culture suspension were collected at the assay endpoint, gently centrifuged (300 × g, 5 min), and resuspended in equal volumes of DPBS. Aliquots were transferred to 96-well plates, where each well was visually divided into defined regions of equal area, and the number of spheroids within each region was counted under a brightfield microscope. Spheroid counts per condition were normalised to the average of the matched control (set to 100%) and expressed as relative spheroid-forming ability (%).

### 3D cell viability assay

Cell viability in 3D spheroid cultures was assessed using the CellTiter-Glo® 3D Cell Viability Assay (Promega, G9682) according to the manufacturer’s instructions. Cells were embedded in 10% methylcellulose supplemented with 1% (v/v) Matrigel (Corning, CLS354234) and seeded at 500 cells per well in 96-well round bottom microplate (Corning) to allow spheroid formation. At the indicated endpoints, plates were equilibrated to room temperature, and an equal volume of CellTiter-Glo 3D reagent was added per well. Plates were placed on an orbital shaker for 5 min to ensure spheroid lysis, followed by 40 min at room temperature for luminescent signal stabilisation. Luminescence was measured using a microplate reader (BioTek Synergy). Values were normalised to the matched control condition and expressed as percentage viability. Three biological replicates were performed per condition.

### RNA isolation, cDNA conversion and quantitative PCR

Total RNA was extracted using TRIzol Reagent (Thermo Fisher, 15596018) according to the manufacturer’s instructions, with isopropanol precipitation. RNA concentration and purity (A260/A280 ≥ 1.9) were assessed using a NanoDrop spectrophotometer (Thermo Fisher). cDNA was synthesised from 1 µg of total RNA using the LunaScript RT SuperMix Kit (NEB, E3010). Quantitative PCR was performed using QuantiNova qPCR Master Mix (Qiagen, 208252) on a LightCycler 480 instrument II (Roche). Reactions were performed in technical triplicate, with three biological replicates per condition. Gene expression was quantified by the comparative C_t_ (ΔΔC_t_) method with β-actin (ACTB) as the internal reference gene. Primer sequences are provided in Supplementary Table S5.

### PCR-based detection of CPE transcript variants

To examine the presence of *CPE* transcripts containing different 5′ regions, a PCR-based detection method was performed on cDNA prepared from 2D and 3D cultures of MCF7 and MCF12A cells of control and ARNT-knockout conditions.

Primer pairs were used (Supplementary Table S6):

***(i)*** CPE_FULL_NTERMINAL_FWD + CPE_NTERMINAL_REV, spanning canonical *CPE* 5′ start codon region (expected product 388 bp);
***(ii)*** CPE_ALTERNATIVE_FORM_4ATG_FWD + CPE_FULLCDNA_REV, starting from the described CPE-ΔN start codon (expected product 952 bp),
***(iii)*** CPE_ALTERNATIVE_FORM_5ATG_FWD + CPE_FULLCDNA_REV, starting from an internal ATG start codon (expected product 760 bp); and
***(iv)*** CPE_ALTERNATIVE_FORM_6ATG_FWD + CPE_FULLCDNA_REV, starting from downstream internal ATG start codon (expected product 700 bp) to distinguish differential amplicons based on intensities.
***(v)*** CPE_CONSERVED_FWD and CPE_CONSERVED_REV primer pair were used for the amplification of a conserved *CPE* coding region, as a reference transcript.

PCR was performed with 150 ng cDNA using Q5® High-Fidelity DNA Polymerase (NEB, M0491) under the following conditions: 98 °C for 30 sec; 35 cycles of 98 °C for 10 s, 59-61 °C for 20 s, 72 °C for 30 s; final extension 72 °C for 5 min. Products were resolved on 2% agarose gels with a 100 bp DNA ladder (NEB, SM0241).

### Whole-cell lysate preparation and immunoblot analysis

Whole-cell lysates were prepared as previously described ^30^ in RIPA buffer (50 mM Tris-HCl pH 7.5, 150 mM NaCl, 1% NP-40, 0.5% sodium deoxycholate, 0.1% SDS) supplemented with cOmplete EDTA-free protease inhibitor cocktail (Roche, 11873580001) and phosphatase inhibitors (Roche, 4906845001). Lysates were cleared by centrifugation at 16,000 × g for 15 min at 4 °C, and protein concentration was determined by Thermo Scientific™ Pierce™ Bradford Plus Protein Assay Kit (Thermo, 10741945). Equal amounts of protein (30 µg per lane) were resolved by SDS–PAGE on Mini-PROTEAN® Precast Gel and transferred to PVDF membranes (Millipore, IPVH00010). Membranes were blocked in 5% (w/v) BSA in TBS-T (TBS, 0.1% Tween-20) for 2 h at room temperature and incubated overnight at 4 °C with primary antibodies as listed below. After three washes in TBS-T, membranes were incubated with HRP-conjugated secondary antibodies (anti-rabbit, Abcam ab97051, RRID:AB_10679369; anti-mouse, Abcam ab97023, RRID:AB_10679675) at 1:5,000 for 1 h at room temperature and visualized using ECL reagent (Bio-Rad, 1705061) on a ChemiDoc imager (Bio-Rad).

Primary antibodies used were: ARNT (Cell Signaling Technology, 5537; 1:1,000); CPE (Proteintech, 86111-1-RR; 1:2,500); Bax (Abcam, ab182734; 1:1,000); Bcl-2 (St John’s Laboratory, STJ27524; 1:1,000); Vinculin (Elabscience, AN005300L; 1:1,000); Histone H3 (Cell Signaling, 9715; 1:1,000); GAPDH (Elabscience, E-AB-40516; 1:1,000).

### Subcellular fractionation

Subcellular fractionation was performed to separate cytoplasmic and nuclear protein fractions. Briefly, cells (∼5 × 10⁶ per condition) were harvested, washed in ice-cold PBS, and resuspended in 200 µl of Buffer A (10 mM HEPES pH 7.9, 1.5 mM MgCl₂, 10 mM KCl, 0.5 mM DTT, 0.5 mM PMSF) supplemented with cOmplete protease inhibitor cocktail. After 10 min on ice, NP-40 was added to a final concentration of 0.1% (v/v), and lysates were vortexed for 10 s and centrifuged at 5,000 × g for 5 min at 4 °C. The supernatant (cytoplasmic fraction) was collected. The nuclear pellet was washed twice with Buffer A, then resuspended in 100 µl of Buffer B (20 mM HEPES pH 7.9, 1.5 mM MgCl₂, 420 mM NaCl, 0.2 mM EDTA, 25% glycerol, 0.5 mM DTT, 0.5 mM PMSF, protease inhibitors). Nuclear lysates were sonicated for 40 cycles as 10 sec ON/10 sec OFF (Bioruptor) to disrupt chromatin and clarified by centrifugation at 16,000 × g for 15 min at 4 °C. Soluble cytoplasmic and nuclear fractions were quantified by Bradford assay and analysed by immunoblot as described. Fractionation purity was assessed using GAPDH (cytoplasmic marker) and Histone H3 (nuclear marker). Three biological replicates were performed.

### RNA-seq

Total RNA was extracted from MCF7 (CNT, ARNT KO, ARNT OE) under matched 2D and 3D conditions using TRIzol. For each culture condition, RNA-seq included four biological replicates for the non-targeting control, three for ARNT knockout and two for ARNT overexpression. RNA integrity was assessed using a Bioanalyzer 2100 (Agilent, G2939BA) — only samples with RIN ≥ 8 were processed. Stranded mRNA libraries were prepared using the Illumina® Stranded mRNA Prep Kit (Illumina, 20040534) and sequenced on an Illumina NovaSeq 6000 instrument (paired-end) to a target depth of 25–30 million reads per sample.

Quality control of raw and trimmed reads was performed using FastQC (version 0.12.1). Raw paired-end RNA-seq reads were adapter- and quality-trimmed using Trim Galore (version 0.6.10) with Cutadapt (version 4.9) in paired-end mode using a Phred quality cutoff of 20. Trimmed reads were aligned to the human hg38 reference genome using STAR ^31^ (version 2.7.10b) with GENCODE v46 annotation, ‘--quantMode TranscriptomeSAM’, and ‘--outSAMtype BAM SortedByCoordinatè, producing coordinate-sorted genomic BAM and transcriptome-aligned BAM files to generate both coordinate-sorted genomic BAM files and transcriptome-aligned BAM files. Transcriptome-aligned BAM files were quantified with RSEM ^32^ (version 1.2.28) using ‘rsem-calculate-expression --alignments --paired-end -p 10’ and the RSEM reference. For each sample, transcriptome-aligned BAM files generated by STAR were supplied to rsem-calculate-expression in paired-end alignment mode, and gene-level expression estimates were obtained from the resulting *.genes.results files. Alignment file inspection and handling were performed with samtools (version 1.6).

Gene-level expected counts generated by RSEM were imported into R (version 4.2.3) using tximport (version 1.26.1) with type = “rsem”. The imported expression matrix included non-targeting control samples in 2D and 3D culture conditions (NT_2D, n = 4; NT_3D, n = 4), ARNT knockout samples in 2D and 3D conditions (ARNT_KO_2D, n = 3; ARNT_KO_3D, n = 3), and ARNT overexpression samples in 2D and 3D conditions (ARNT_OE_2D, n = 2; ARNT_OE_3D, n = 2). DESeq2 ^33^ (version 1.38.3) was used to construct count-based expression models and perform differential expression analysis and the results were extracted with ‘independentFiltering = FALSÈ. Differential expression testing was conducted using a design formula based on experimental condition, enabling comparisons between ARNT knockout, ARNT overexpression and non-targeting control groups within matched 2D and 3D culture contexts, as well as comparisons between 2D and 3D conditions within each experimental group. Statistical output included normalized expression estimates, log_2_ fold-change values, Wald test statistics, raw p-values, and multiple-testing adjusted p-values.

For gene-level annotation, Ensembl identifiers were processed to remove version suffixes before mapping to gene symbols. Annotation was performed in R using AnnotationDbi (version 1.60.2) and org.Hs.eg.db (version 3.16.0). Annotated differential expression tables were then exported for downstream interpretation and visualization.

Principal component analysis was performed for exploratory sample-level quality assessment using variance-stabilized expression values generated with DESeq2::vst. PCA was calculated using the top 500 most variable genes after filtering lowly expressed genes. Pairwise PCA visualizations were generated by subsetting the same global PCA coordinate space for selected sample-group comparisons.

A focused response-pattern heatmap was generated using a custom Python 3.12.13 workflow implemented with NumPy 2.0.2, pandas 2.2.2, Matplotlib 3.10.0 and Seaborn 0.13.2. Gene-level differential expression results were obtained from filtered DESeq2 output tables derived from ARNT perturbation contrasts in 2D and 3D culture conditions, including KO versus NT (2D), OE versus NT (2D), KO versus NT (3D), OE versus NT (3D) and KO versus OE (3D). A predefined set of biologically relevant genes (*ARNT, LXN, CPE, CDKN1A, GSN, SLIT1, TRIM29, ACTA2, AHRR* and *SYT7*) was selected, and the corresponding log_2_ fold-change values were compiled into a summary matrix for comparative visualization.

To enable biologically informed interpretation, genes were grouped according to response patterns observed across ARNT loss- and gain-of-function conditions. Genes showing positive regulation in KO versus NT (3D) together with negative regulation in OE versus NT (3D) were classified as reciprocal/opposite-response, reflecting inverse behaviour under ARNT depletion and overexpression in 3D culture. Genes detected in the filtered KO versus NT (3D) results but not in the corresponding filtered KO versus NT (2D) results were classified as 3D-specific responses, consistent with context-dependent regulation. Genes showing negative fold-change in both KO versus NT (2D) and KO versus NT (3D) were classified as core ARNT-dependent, whereas genes showing positive fold-change in both knockout contrasts were classified as shared KO response. Genes were ordered by response class before visualization, and the resulting matrix was displayed as a heatmap with a zero-centred diverging colour scale. Missing values were masked and shown in grey, and response-group structure was further highlighted with bracket annotations and separator lines.

### CCLE ABSOLUTE copy-number analysis

Allele-specific copy-number profiles for breast cancer cell lines were obtained from the Cancer Cell Line Encyclopedia (CCLE) ABSOLUTE call set (CCLE_ABSOLUTE_combined_20181227, downloaded from the DepMap portal). The dataset comprises segmented total and allele-specific copy-number calls together with sample-level purity, ploidy and inferred genome-doubling status for 52 breast cancer cell lines. For each sample, segments overlapping the q-arm of chromosome 1 (defined as Chr1 coordinates ≥ 125 Mb on the hg19 reference) were extracted. For each cell line, length-weighted mean total copy number, median total copy number, and the percentage of Chr1q with total copy number above and below sample ploidy were computed. Cell lines were classified as ‘broad gain’ (≥50% of Chr1q above ploidy), ‘partial gain’ (20–50%), ‘broad loss’ (≥50% below ploidy), ‘partial loss’ (20–50%) or ‘no major change’ otherwise. Per-cell-line copy-number profiles for MCF7 and T47D are shown in (Supplementary Fig. S1A), and the full panel summary is provided in Supplementary Table S7. Analysis was performed in Python (version 3.10) using pandas, NumPy and matplotlib.

### Public ChIP–seq data and motif analysis

Public ChIP-seq peak collections for ARNT and ESR1 were retrieved from ChIP-Atlas^34,35^ using the Peak Browser interface with the genome assembly set to hg38, the track type class set to “ChIP: TFs and others”, the cell type class restricted to “Breast”, the significance threshold set to 50, and the track types specified as ARNT and ESR1. The resulting collections were then downloaded in BED format. The CPE locus was used as the query region in a custom Python workflow implemented in Python 3.12 with requests 2.32.4, NumPy 2.0.2 and Matplotlib 3.10.0. Genomic coordinates for CPE were resolved programmatically, and the analysis window was defined by extending the locus by 120 kb upstream and downstream. Peak intervals overlapping the selected region were extracted for each factor and aggregated into 500 equally spaced genomic bins to generate locus-level support profiles, in which support was defined as the number of overlapping peak intervals per bin. The distribution of individual peak calls was additionally represented by peak-center tick marks along the genomic axis. Gene annotations overlapping the queried interval were retrieved programmatically and incorporated into the final multi-panel genomic plot, which displayed the gene model together with ARNT and ESR1 support tracks across the CPE locus.

### Motif and proximity analysis of ARNT-bound CPE locus peaks

ARNT peaks overlapping the CPE ±120 kb locus were used for sequence-level motif analysis. Custom Python workflows were implemented in Python 3.12 with NumPy 2.0.2, pandas 2.2.2, Matplotlib 3.10.0, requests 2.32.4 and SciPy 1.16.3. Peak sequences were retrieved from the hg38 reference genome and scanned using both position-weight-matrix-based and exact-match approaches. JASPAR 2026 ^36^ vertebrate motif models corresponding to Arnt, Ahr::Arnt and ARNT::HIF1A were evaluated using FIMO v5.5.9 ^37^, whereas exact sequence searches were performed for the canonical E-box sequence CACGTG and the shorter E-box core sequences CGTG and CACG. The same ARNT peak sequences were additionally scanned using the JASPAR ESR1 motif model and exact searches for the full estrogen response element pattern GGTCAnnnTGACC and the half-ERE sequence GGTCA, including reverse-complement half-site occurrences. FIMO analyses were performed using a significance threshold of p<1×10⁻⁴. Exact motif occurrences were converted from peak-relative positions to hg38 genomic coordinates and summarized at the ARNT peak level, together with peak coordinates, peak score, location relative to the CPE gene body, distance to the CPE transcriptional start region and motif occurrence counts. ESR1 intervals overlapping the CPE locus were sorted and merged using bedtools v2.27.1^38^ to generate nonredundant ESR1 regions, and each ARNT peak was assigned to its nearest merged ESR1 region using BED-style interval distance and center-to-center offset. These motif and proximity annotations were incorporated at locus-level to indicate E-box/core motif-positive and half-ERE-positive ARNT peaks, their positional relationship to ESR1-supported regions and aggregate motif counts across the ARNT peak set.

### ESR1–CPE expression correlation analysis

Spearman correlation between *ESR1* and *CPE* expression was computed in Luminal A samples from METABRIC (n = 700; log2-transformed Illumina microarray values) and TCGA-BRCA PanCancer Atlas (n = 420; log2(RSEM+1)-transformed RNA-seq values). Two-sided p-values are reported throughout. METABRIC expression data were obtained from cBioPortal ^39^.

### TCGA-BRCA Data Integration and Hypoxia Enrichment Analysis

Molecular and clinical datasets for the TCGA-BRCA cohort were retrieved from the UCSC Xena Browser ^40^. Transcriptomic profiles were obtained as RSEM-normalized RNA-seq data, while clinical annotations, including PAM50 subtypes and ER immunohistochemistry status, were integrated using patient-level identifiers. To maintain data consistency, TCGA barcodes were truncated to 12-character patient IDs, and in instances where multiple entries existed for a single patient, expression and clinical values including ABSOLUTE tumour purity and ploidy estimates were averaged. For downstream analyses, log2(RSEM+1) transformation was applied to the expression levels of ARNT and CPE. To quantify hypoxia-related transcriptional activity, single-sample gene set enrichment analysis (ssGSEA^41^) was performed via the gseapy 1.2.1 library using the 200-gene HALLMARK_HYPOXIA set from MSigDB ^42^ (of which 191 genes were present in the filtered matrix). This process yielded a final, quality-controlled analytical cohort of 1,082 samples, each containing complete records for ARNT/CPE expression, hypoxia enrichment scores, and corresponding clinico-pathological metadata, ensuring the removal of all duplicate entries and invalid identifiers.

### Single-cell RNA sequencing analysis

Breast cancer single-cell RNA-seq data from Wu et al. 2021 (GSE176078) ^43^ were analyzed using Google Colab in Python. Data was downloaded from the GEO database. The processed count matrix, gene annotation, barcode, and metadata files were imported using the Scanpy (Version 1.12.1) /AnnData (Version 0.12.14) framework. An AnnData object was generated by combining the count matrix with the corresponding gene and cell barcode information.

Cell-level metadata, including sample identity, breast cancer subtype, and cell type annotations, were integrated into the dataset using cell barcodes. Only cells annotated as “Cancer Epithelial” were selected for downstream analysis in order to focus on tumor cells.

Expression counts were normalized to 10,000 counts per cell and log-transformed using Scanpy normalization functions. Log-normalized expression values of ARNT and CPE, together with sample and subtype metadata, were extracted and exported as a CSV file for further analysis.

Subsequent statistical analyses were performed in Python using pandas (Version 2.3.3), numpy (Version 2.0.2), scipy.stats (Version 1.16.3), and matplotlib (Version 3.10.0). Cancer epithelial cells with detectable CPE expression were defined as cells with log-normalized CPE expression values greater than zero (CPE > 0). The proportion of CPE-expressing cells was calculated for each molecular subtype (ER+, TNBC, and HER2+), and differences between groups were evaluated using the Mann– Whitney U test and visualized using bar plots.

For within-subtype analysis, ER+ cancer epithelial cells with detectable ARNT expression (ARNT > 0, n = 1,410) were selected. Cells were clustered into ARNT-high and ARNT-low groups using the median ARNT expression value as the threshold. Differences in CPE expression between groups were assessed using the Mann– Whitney U test. Data visualization was performed using matplotlib in Python.

### Statistical analysis

All quantitative experiments were performed in three independent biological replicates unless otherwise specified, with technical triplicates within each biological replicate where applicable. Data are presented as the mean ± standard error of the mean (s.e.m.) unless otherwise indicated. For comparisons between two groups, two-tailed Student’s *t*-tests were used; Welch’s correction was applied where variances were unequal (F-test *P* < 0.05). For non-normally distributed data (assessed by D’Agostino– Pearson omnibus test), the Mann–Whitney U test or Kruskal–Wallis test with Dunn’s post-hoc were used. For spheroid area measurements, comparisons between groups were performed using the Student t-test following log-transformation to approximate normality. Significance levels are denoted as: ns, *\*P* < 0.05; \*\**P*< 0.01; \*\*\**P* < 0.001; \*\*\*\**P* < 0.0001. All *P*-values are two-sided. Statistical analyses were performed using GraphPad Prism (version 10.0). Sample sizes (n) reflect biological replicates and are reported in each figure legend.

For high-throughput data (CRISPR screen, RNA-seq), false discovery rates (FDR) were computed by Benjamini–Hochberg correction. CRISPR screen significance thresholds were set at FDR < 0.1 (single-condition dropout) and FDR < 0.25 (3D vs 2D enrichment); RNA-seq DEG significance was set at FDR < 0.05. No statistical methods were used to predetermine sample sizes; sample sizes were chosen based on standard practice in the field for the corresponding assays. No data were excluded from analyses. Investigators were not blinded during data acquisition or analysis, given the genotype-driven nature of the experimental conditions.

### Reproducibility

All experiments were performed with at least three independent biological replicates unless otherwise stated. The number of replicates and the statistical test applied are reported in each figure legend. Representative western blot images shown in figures are from one of three independent experiments yielding consistent results. Brightfield spheroid images were selected as representative of the typical appearance from at least 50-60 spheroids imaged per replicate across three biological replicates.

## Results

### Parallel CRISPR screens in 2D and 3D cultures identify ARNT as an architecture-dependent fitness gene

To identify chromosome 1q-encoded genes whose contribution to cell fitness depends on culture architecture, we performed parallel CRISPR–Cas9 loss-of-function screens in matched two-dimensional (2D) monolayers and three-dimensional (3D) spheroids of the MCF7 and T47D luminal breast cancer cell lines. Analysis of CCLE ABSOLUTE data ^44^ confirmed that Chr1q gain is common among breast cancer cell lines, with 48 of 52 lines (92%) carrying broad or partial Chr1q gains (Supplementary Fig. S1A; Supplementary Table S7). We selected the MCF7 cell line, which harbours focal 1q gains on a near-diploid background (ploidy 2.02), and the T47D cell line, which shows broad 1q amplification on a genome-doubled background (ploidy 2.89), as complementary genomic contexts. We generated a custom sgRNA library, LIBGENE1Q, targeting 825 non-essential Chr1q-encoded genes, together with sgRNAs targeting 80 common essential genes (positive controls), 2 additional external positive-control genes, and 500 non-targeting controls (Supplementary Fig. S1B; Supplementary Table S1).

Cells were transduced at a MOI below 0.4 to ensure single sgRNA integration, and library coverage was maintained above 1,000-fold throughout the 20-day screening period (Fig. 1A). For each cell line, parallel cultures were maintained as 2D monolayers and 3D spheroids to enable direct comparison of fitness dependencies across culture conditions. Library representation and inter-replicate reproducibility were confirmed by analysis of sgRNA abundance distributions across time points and replicates (Fig. 1B; Supplementary Fig. S1C-E). Screen performance was further assessed by comparing the depletion of sgRNAs targeting common essential genes (CE) with behaviour of non-targeting controls (NT). CE-targeting sgRNAs were significantly depleted at the endpoint in both culture conditions, while NT sgRNAs remained stable (Fig. 1C). The magnitude of CE depletion was lower in 3D than in 2D, in line with the previously reported attenuation of essential gene phenotypes in spheroid culture^11^. MAGeCK-RRA analysis^28^ identified 18 non-essential genes that met the depletion criteria in both MCF7 and T47D cells (FDR < 0.1, log_2_FC < −0.8) (Supplementary Fig. S1F; Supplementary Table S8A-D).

**Fig. 1.**
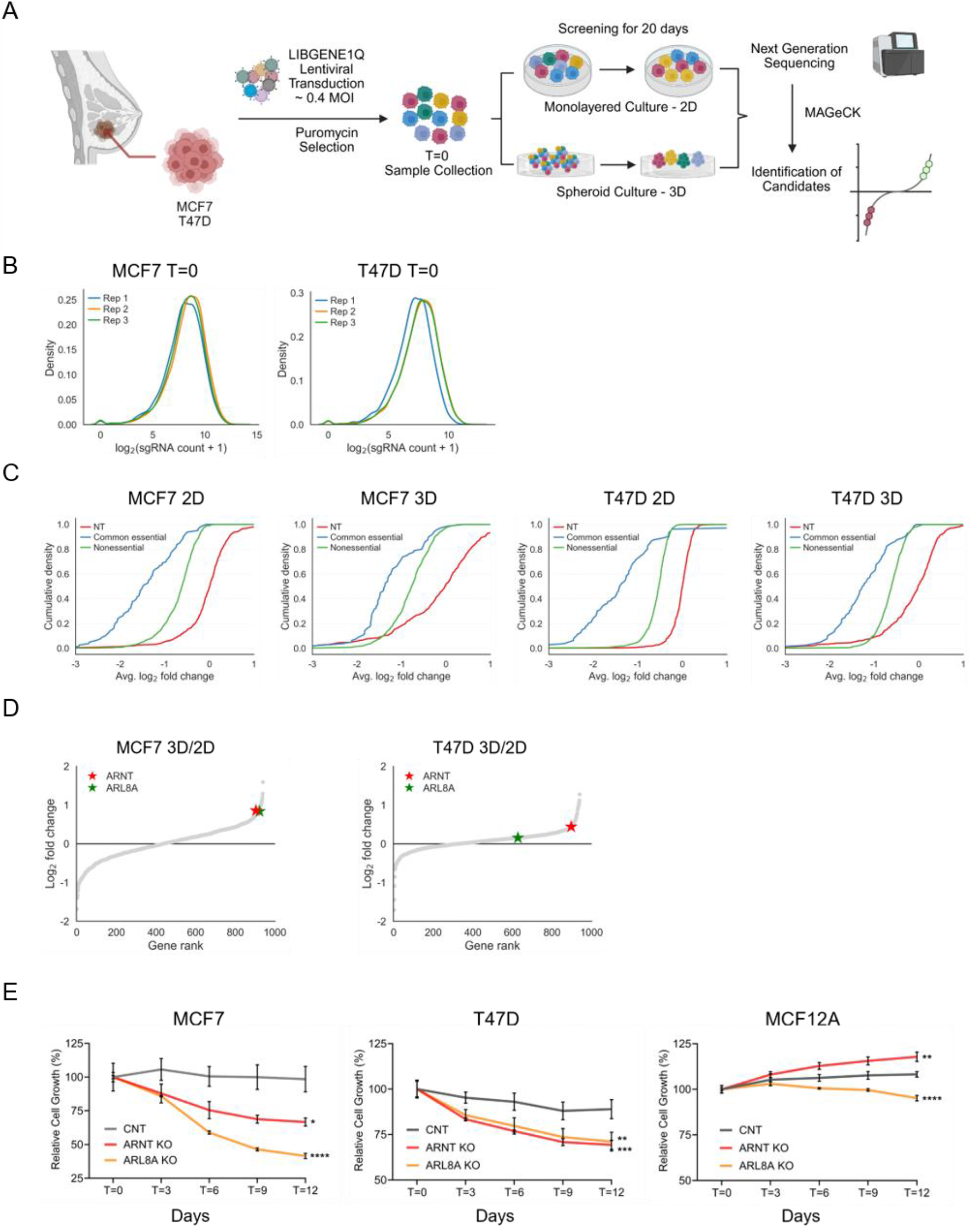
Parallel CRISPR screening in 2D and 3D cultures identifies architecture-dependent fitness genes in breast cancer. **A** Schematic of the pooled CRISPR screening workflow in MCF7 and T47D cells using the LIBGENE1Q library under parallel 2D monolayer and 3D spheroid conditions. **B** Density plots of sgRNA counts at T=0 across three biological replicates in MCF7 and T47D. **C** Cumulative distribution plots of log_2_ fold change (T=20 vs. T=0) for common essential (CE), non-essential (NE) and non-targeting (NT) sgRNAs in MCF7 and T47D under 2D and 3D conditions. **D** Gene rank plots of 3D/2D-normalized MAGeCK-RRA scores in MCF7 and T47D; *ARNT* and *ARL8A* are highlighted. **E** Competitive growth assays over 12 days in MCF7, T47D and MCF12A. Data represent mean ± s.e.m. from three biological replicates. *P<0.05 (*), P<0.01 (**), P<0.001 (***)* and *P<0.0001 (****)* by two-tailed Student’s *t*-test.

To identify genes with differential fitness effects between 2D and 3D conditions, we normalised the 3D sgRNA enrichment scores to their corresponding 2D scores, generating a 3D/2D ratio. Of the 109 genes depleted in MCF7 cells (T=20 and T=0), 8 showed an enrichment profile (log_2_FC > 0.75), compared with 4 enriched genes in T47D cells (Supplementary Table S8E, F). We therefore used MCF7 as the discovery set for architecture-dependent candidates and retained T47D as an independent model for validation. Among the MCF7 candidates, *ARNT*-and *ARL8A*-targeting sgRNAs were depleted in both culture conditions, but their depletion was weaker in 3D, resulting in relative enrichment in the 3D/2D comparison (Fig. 1D; Supplementary Fig. S1G). ARNT shows a broad mutational landscape across diverse cancer types ^45^ and has been reported to exert both tumour-promoting and tumour-suppressive effects, making it a compelling candidate for architecture-dependent regulation.

To independently validate the screen-derived fitness effects of ARNT and ARL8A, we performed competitive growth assays to test whether sgRNA-mediated targeting of either gene altered cell fitness. GFP-labelled reference cells were mixed with sgRNA-transduced cells targeting ARNT or ARL8A at a 30:70 ratio, and changes in population composition (GFP proportion) were tracked by flow cytometry over 12 days as a readout of relative fitness ^29,46^. An increase in the GFP-positive reference population therefore indicated reduced fitness of the gene-targeted cells. In both MCF7 and T47D cells, targeting of *ARNT* or *ARL8A* reduced fitness in 2D, consistent with their depletion in the primary screen (Fig. 1E; Supplementary Fig. S1H). In MCF7 cells, ARL8A loss conferred a stronger fitness disadvantage than ARNT loss. By contrast, *ARNT* loss produced a slight increase in relative fitness in the non-tumorigenic MCF12A mammary epithelial cells, whereas *ARL8A* loss had only a marginal negative effect (Fig. 1E; Supplementary Fig. S1H). Thus, the ARNT fitness requirement observed in 2D luminal breast cancer cells was not reproduced in MCF12A cells. The reproducibility of the ARNT phenotype across two independent luminal breast cancer models, its differential behaviour between 2D and 3D screens, and the absence of a comparable fitness defect in MCF12A cells led us to prioritise ARNT for detailed functional characterisation.

### ARNT exerts opposing fitness effects in 2D and 3D cultures

To determine whether tumour architecture influences ARNT-dependent fitness, we systematically assessed the effects of *ARNT* loss and overexpression in 2D monolayer and 3D spheroid cultures of MCF7, T47D and MCF12A cells. ARNT depletion and overexpression were confirmed at both protein and mRNA levels prior to functional analysis; complete loss of ARNT protein by immunoblot confirmed effective depletion in each model (Supplementary Fig. S2A, B).

In 2D colony formation assays, *ARNT* knockout reduced clonogenic capacity in MCF7 and T47D cells consistent with the differential fitness effects observed in the primary screen, while MCF12A cells exhibited enhanced clonogenic growth upon *ARNT* loss. Conversely, ectopic ARNT overexpression strongly increased colony formation in MCF7 cells, produced minimal effects in T47D cells and reduced clonogenic growth in MCF12A cells (Fig. 2A, B).

**Fig. 2.**
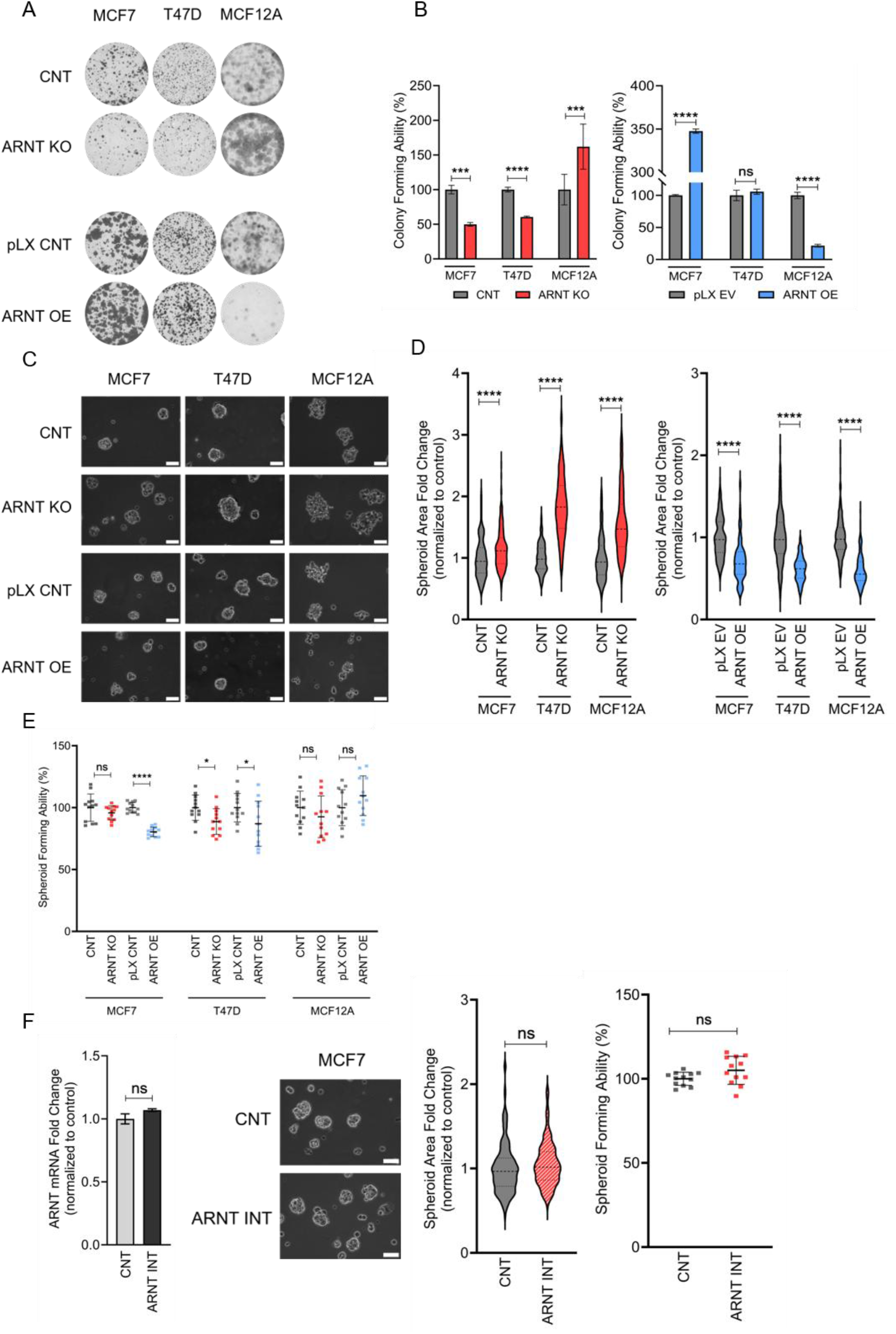
ARNT modulation produces inverted fitness phenotypes in 2D and 3D breast cancer models. **A** Representative image of colony formation assays in MCF7, T47D and MCF12A cells following ARNT knockout (ARNT KO) or overexpression (ARNT OE) and their respective controls. **B** Quantification of colony-forming ability under 2D conditions. **C** Representative brightfield images of 3D spheroids; scale bar, 70 µm. **D** Quantification of spheroid area fold change following ARNT KO (left) and ARNT OE (right). **E** Spheroid-forming ability across cell lines. **F** Validation of phenotype specificity using an intergenic sgRNA control (ARNT-INT) targeting a region proximal to the ARNT locus without disrupting ARNT expression: ARNT mRNA expression levels normalized to Beta-actin and presented relative to control (left), representative brightfield images of spheroids formed following intergenic targeting (center), spheroid area fold change normalized to control (right), and spheroid-forming ability (far right) in MCF7 cells. Data represent mean ± s.e.m. from three biological replicates. *P<0.05 (*), P<0.01 (**), P<0.001 (***)* and *P<0.0001 (****)*; ns, not significant; two-tailed Student’s *t*-test.

We next assessed the effect of ARNT perturbation on 3D spheroid growth. In contrast to its growth-inhibitory effect in 2D luminal breast cancer cells, *ARNT* knockout increased spheroid area in MCF7, T47D and MCF12A cells, indicating that *ARNT* loss promotes spheroid expansion across the tested models. ARNT overexpression produced the opposite effect, reducing spheroid growth in all three cell lines and yielding distorted, non-compact spheroid structures in MCF12A cells (Fig. 2C, D). Spheroid-forming ability assays confirmed that ARNT overexpression reduced spheroid formation frequency in MCF7 and T47D cells, while *ARNT* loss did not significantly alter spheroid formation frequency despite increasing spheroid size (Fig. 2E). Longitudinal quantification by CellTiter-Glo 3D further showed a progressive increase in viable cell mass following *ARNT* targeting in MCF7 spheroids (Supplementary Fig. S2C). In line with these findings, ARNT loss in MCF12A spheroids was associated with up-regulation of the pro-survival protein Bcl-2, whereas ARNT overexpression was accompanied by increased Bax expression, consistent with altered survival signalling in the spheroid context (Supplementary Fig. S2D).

To examine whether these phenotypes reflect ARNT-specific activity rather than regional effects associated with Chr1q copy number, we targeted a non-coding intergenic region proximal to the *ARNT* locus using a sgRNA (ARNT-INT) that did not alter ARNT expression. Intergenic targeting did not significantly affect spheroid area or spheroid-forming ability in MCF7 cells (Fig. 2F), supporting the conclusion that the spheroid phenotype is attributable to *ARNT* gene function rather than to regional copy-number-associated targeting effects. We further validated phenotype specificity in rescue experiments using a PAM-mutant ARNT overexpression construct that restored *ARNT* expression in knockout cells. Re-expression of PAM-mutant ARNT reversed the increased spheroid growth induced by *ARNT* loss (Supplementary Fig. S2E-H).

Together, these results establish that ARNT supports fitness in 2D luminal breast cancer cells but restricts spheroid growth, revealing opposing architecture-dependent effects consistent with the dimensionality-aware enrichment pattern identified in the primary screens.

### CPE had been characterized as a downstream effector of ARNT in 3D spheroids

To determine how culture architecture reshapes the ARNT-dependent transcriptional programme, we performed transcriptomic profiling of matched 2D monolayer and 3D spheroid cultures of MCF7 cells with stable ARNT knockout (KO) or overexpression (OE), each compared with the corresponding control cells (CNT) (Supplementary Table S9). Differential expression analysis using DESeq2 ^33^ confirmed successful *ARNT* perturbation across all conditions (KO: log_2_FC ≈ −1.28; OE: log_2_FC = +1.26 to +2.08; *P*-adjusted < 10^−16^). The number of differentially expressed genes (DEGs) was substantially higher in 2D than in 3D cultures following both perturbations (KO: 144 vs 18 DEGs; OE: 216 vs 34 DEGs), indicating that the transcriptional response to ARNT modulation is more restricted in the spheroid context. In 2D, ARNT KO resulted in upregulation of genes associated with stress-response and cell-cycle arrest programmes (e.g. *CDKN1A*, *BTG2*, *NUPR1*, *DDIT3*) and downregulation of survival-associated genes (e.g. *TERT*, *SLC7A5*, *NFE2L2*). These data were consistent with the reduced fitness observed following ARNT loss in monolayer culture. Conversely, ARNT OE in 2D resulted in significant downregulation of genes associated with differentiation and IGF-sequestering programmes (e.g. *PPARG*, *IGFBP3*, *CEACAM5/6*), consistent with the enhanced colony-forming phenotype (Supplementary Fig. S3A).

In contrast to the broad transcriptional response observed in 2D, ARNT perturbation in 3D elicited a markedly attenuated and more selective transcriptional programme.

Using the 2D transcriptome as a reference, comparative analysis identified both shared and context-specific responses (Fig. 3A). Among the most prominent candidates, *CPE* (carboxypeptidase E) ^25^ and *LXN* (latexin) ^47^ showed reciprocal regulation upon ARNT modulation specifically in 3D. *CPE* expression increased following ARNT KO and decreased upon ARNT OE, whereas *LXN* exhibited the opposite pattern (Fig. 3A).

**Fig. 3.**
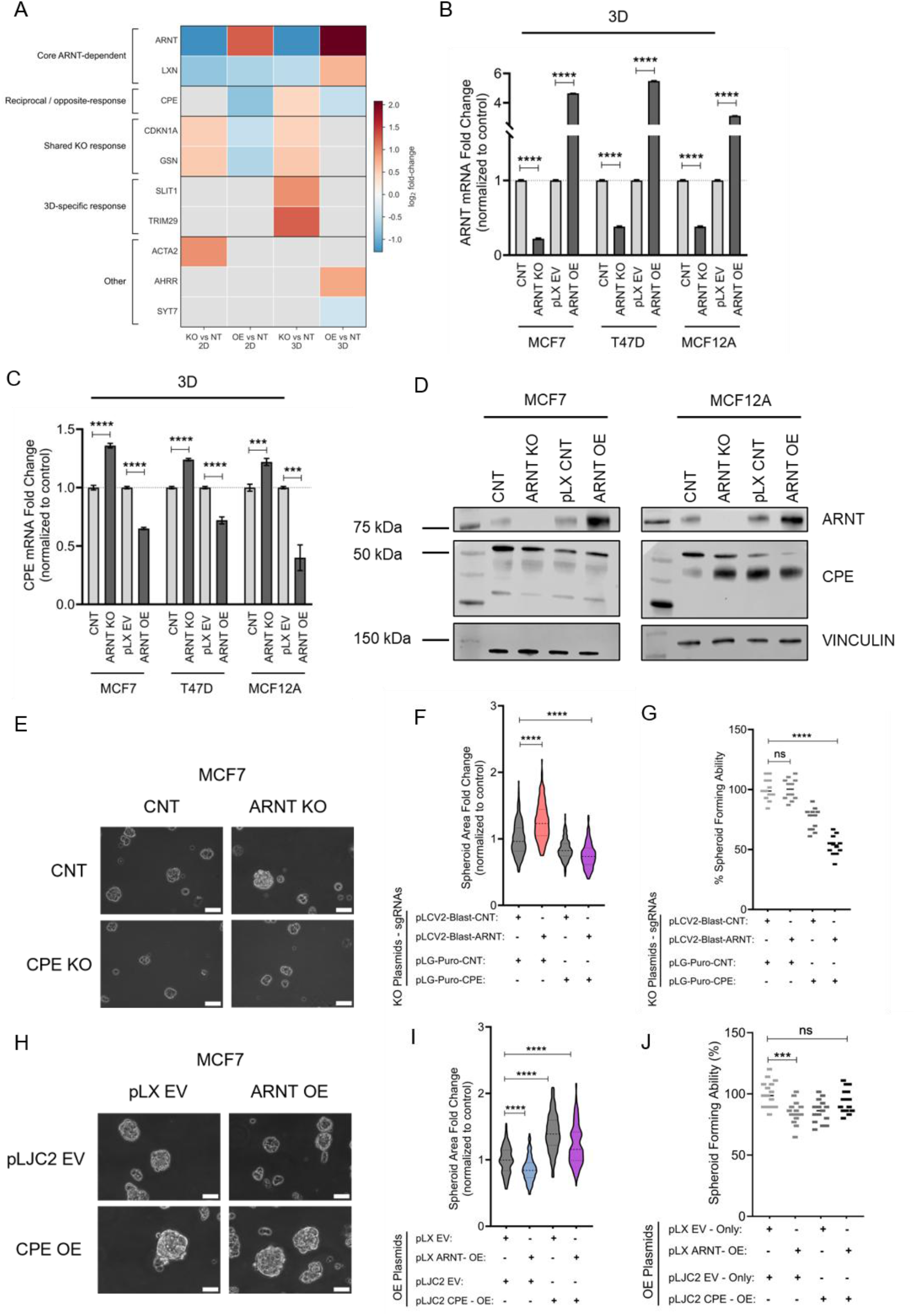
ARNT regulates CPE expression to control 3D spheroid growth in breast cancer. **A** Heatmap of representative differentially expressed genes grouped by directional behavior across four RNA-seq contrasts in MCF7 cells (KO vs. NT 2D, OE vs. NT 2D, KO vs. NT 3D, OE vs. NT 3D); color intensity reflects log_2_fold change. **B** RT-qPCR analysis of ARNT expression in MCF7, T47D and MCF12A spheroids under ARNT KO and ARNT OE conditions, confirming ARNT perturbation status. Expression levels were normalized to Beta-actin and presented relative to the corresponding control group. **C** RT-qPCR validation of CPE mRNA expression in 3D spheroids following ARNT modulation. ARNT loss increased CPE expression whereas ARNT overexpression reduced it, consistent with RNA-seq findings. Expression levels were normalized to Beta-actin and presented relative to the corresponding control group. **D** Immunoblot analysis of ARNT and CPE protein levels in MCF7 (left) and MCF12A (right) spheroids; vinculin, loading control. Molecular weight markers are shown in the leftmost lane of each gel. **E** Representative brightfield images of MCF7 spheroids under double-knockout conditions; scale bar, 70 µm. **F** Spheroid area fold changes under double-knockout conditions. **G** Spheroid-forming ability under double-knockout conditions. **H** Representative brightfield images of MCF7 spheroids under double-overexpression conditions; scale bar, 70 µm. **I** Spheroid area fold change under double-overexpression conditions. **J** Spheroid-forming ability under double-overexpression conditions. Data represent mean ± s.e.m. from three biological replicates. *P<0.05 (*), P<0.01 (**), P<0.001 (***)* and *P<0.0001 (****)*; ns, not significant; two-tailed Student’s *t*-test.

We confirmed this reciprocal regulation at the transcript level by RT-qPCR. *CPE* mRNA expression was increased following ARNT KO and reduced upon ARNT OE in spheroids derived from MCF7, T47D and MCF12A cells (Fig. 3B, C). This regulation was further supported at the protein level. Consistent with the known molecular heterogeneity of CPE expression ^26^, immunoblot analysis of MCF7 detected multiple CPE-immunoreactive species. ARNT loss increased the abundance of several of these species, whereas ARNT overexpression reduced the abundance of full-length CPE (∼53–55 kDa) in MCF7 spheroids and, to a variable extent, in MCF12A spheroids (Fig. 3D).

To assess the functional contribution of CPE and LXN to ARNT-dependent spheroid growth, we independently targeted each gene in MCF7 cells. CPE loss reduced spheroid growth, whereas LXN knockout increased spheroid area (Supplementary Fig. S3B, C), indicating that *CPE* and *LXN* exert opposing effects on spheroid expansion. Further analysis revealed that *LXN* carries a deleterious mutation in T47D cells (Supplementary Fig. S3D), and RT-qPCR confirmed the absence of detectable *LXN* transcript in this cell line. Because *LXN* cannot function as a shared effector across both luminal breast cancer cell lines, we focused subsequent analyses on *CPE*.

To establish epistasis between *ARNT* and *CPE*, we generated double-perturbation models in MCF7 spheroids. *ARNT* KO alone produced spheroid enlargement, whereas simultaneous knockout of *ARNT* and *CPE* reversed this effect and resulted in spheroid shrinkage, demonstrating that CPE is required for the growth-promoting effect of ARNT loss (Fig. 3E-G; Supplementary Fig. S4A). Conversely, ARNT OE reduced spheroid growth, while co-overexpression of *ARNT* and *CPE* restored spheroid growth and spheroid-forming capacity to levels comparable with those of control cells (Fig. 3H-J; Supplementary Fig. S4B). Combined *ARNT* and *CPE* knockout similarly suppressed ARNT-loss-induced spheroid expansion in T47D and MCF12A cells (Supplementary Fig. S4), indicating a conserved requirement for CPE across the tested cellular backgrounds. Together, these findings identify CPE as a functional downstream effector of ARNT and establish a functional ARNT–CPE axis that controls spheroid growth.

### ARNT is associated with the CPE locus and regulates CPE protein distribution in 3D spheroids

Having established CPE as a downstream effector of ARNT in 3D spheroids (Fig. 3E-J; Supplementary Fig. S4), we next examined whether publicly available ChIP–seq datasets supported ARNT occupancy at the CPE locus ^34,35^. Analysis of breast cancer datasets compiled in ChIP-Atlas identified six ARNT ChIP–seq peaks across the CPE genomic locus (Fig. 4A). Motif analysis of these peak sequences revealed the absence of canonical E-box motifs (CACGTG), while a single half-oestrogen response element (half-ERE; GGTCA) was identified at P1 — the peak most proximal to the CPE transcription start site. Given the reported role of ARNT as a coactivator of ERα in luminal breast cancer ^22,23^, we examined the spatial relationship between ARNT and ESR1 occupancy at this locus. All six ARNT peaks were located within 1 kb of ESR1-occupied chromatin regions, and five directly overlapped ESR1 peaks (Fig. 4B-D). These spatial relationships are consistent with the possibility that ARNT is recruited to the CPE locus in association with ERα-bound chromatin, although they do not establish the mechanism of ARNT recruitment.

**Fig. 4.**
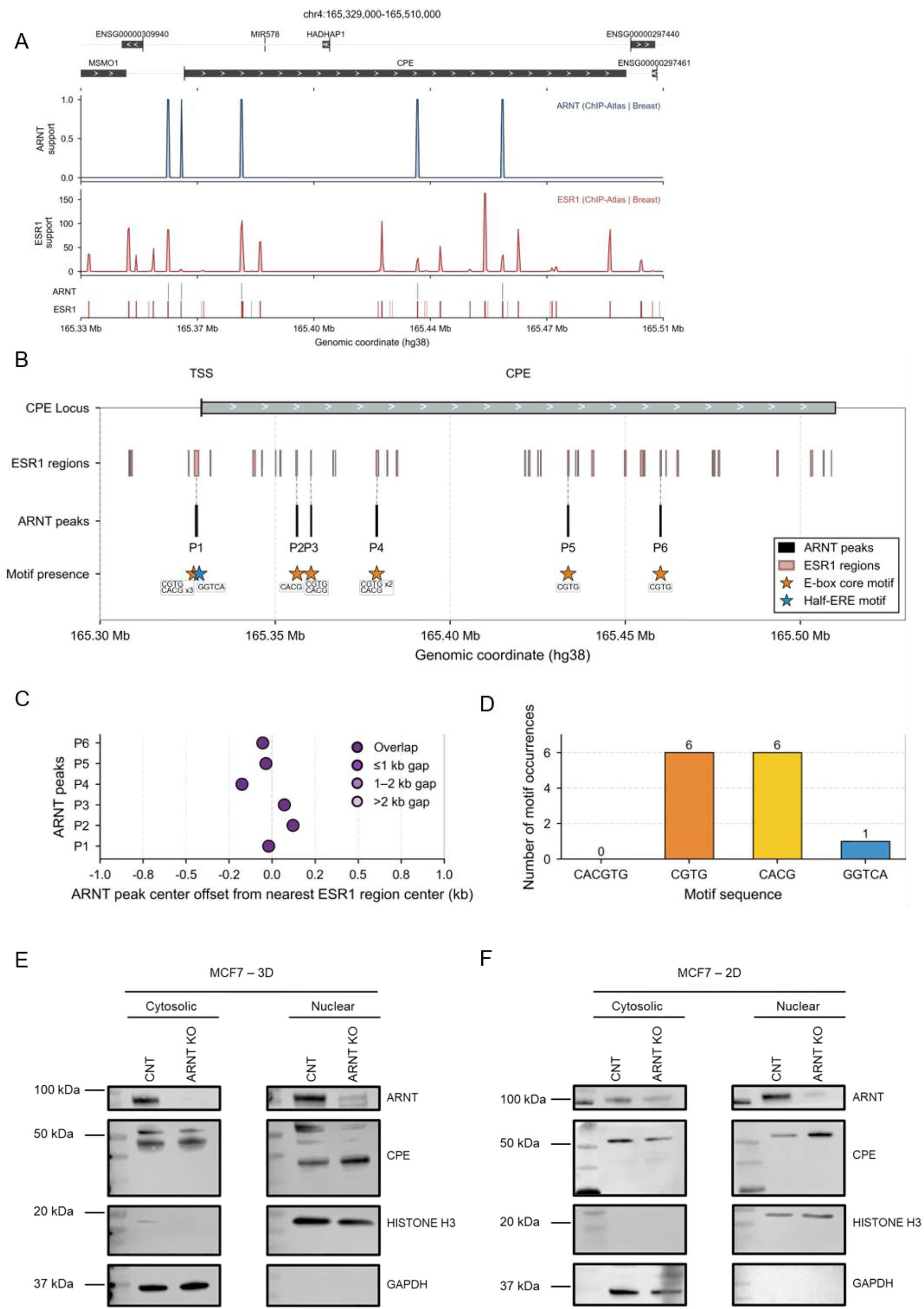
ARNT engages the CPE locus and regulates architecture-dependent CPE protein distribution. **A** Public ChIP-Atlas peak collections at the *CPE* locus (chr4:165,329,000–165,510,000) showing ARNT (top) and ESR1 (bottom) binding profiles in breast tissue. **B** Genomic overview of the *CPE* locus (hg38, chr4: 165.28– 165.52 Mb) showing the positions of ESR1-occupied chromatin regions (pink bars; merged ChIP-Atlas peak collections from breast cancer datasets), ARNT ChIP-seq peaks (black bars; P1–P6), and motif occurrences within peak sequences: E-box core motifs (orange stars; CGTG and CACG) and half-ERE motif (blue star; GGTCA). TSS, transcription start site. **C** ARNT peak center offset from the nearest ESR1 region center (kb) for each peak (P1-P6). Five of six peaks show direct overlap or offset within 0.1 kb; P4 shows an offset of approximately −0.4 kb. **D** Number of motif occurrences across all six ARNT peak sequences. Canonical E-box (CACGTG) was not detected. Partial core sequences CGTG (n = 6) and CACG (n = 6) were identified. A single half-ERE (GGTCA) was detected at P1, the peak most proximal to the CPE TSS. Full ERE (GGTCANNNTGACC) and ESR1/ERα PWM matches (JASPAR 2026, MA0112.4, p < 1 × 10⁻⁴) were absent across all peaks. **E** Cellular fractionation followed by immunoblot analysis of MCF7 cells under control and ARNT KO conditions in 3D spheroid culture. CPE-immunoreactive species at ∼50–53 kDa, ∼40 kDa and ∼27 kDa were detected. Histone H3 and GAPDH were used as nuclear and cytoplasmic loading controls, respectively. **F** Lower molecular weight CPE species (∼27 kDa) were not detected in MCF7 ARNT knockout cells under 2D conditions but became apparent in 3D spheroid cultures only. Molecular weight markers are shown in the leftmost lane of each gel.

To characterise the relationship between ARNT and CPE at the protein level, we performed subcellular fractionation followed by immunoblot analysis using an antibody recognising amino acids 26–453 of CPE. CPE-immunoreactive species showed distinct subcellular distributions in MCF7 3D spheroids (Fig. 4E). A ∼50–53 kDa species corresponding to full-length CPE was detected predominantly in the cytoplasmic fraction, while a ∼40 kDa species consistent with the previously described N-terminal truncated CPE-ΔN isoform was enriched in the nuclear fraction. CPE-ΔN has been reported to arise through translation initiation at the third in-frame ATG of the CPE coding sequence and has been cloned from human cancer cells ^26^. Notably, a ∼27 kDa CPE-immunoreactive species was detected selectively in MCF7 spheroids and was further enriched upon ARNT loss. This species was not detected in MCF7 monolayer cultures or in MCF12A cells, suggesting that its accumulation depends on both culture architecture and cellular context (Fig. 4F; Supplementary Fig. S5A).

To investigate whether lower-molecular-weight CPE species could arise from alternative transcript usage, we designed primer pairs targeting distinct regions of the CPE coding sequence, including sites downstream of the canonical start site (Supplementary Fig. S5B; Supplementary Table S6). Amplicons generated by downstream primer pairs were selectively enriched in MCF7 spheroids relative to monolayer cultures and further increased upon ARNT loss (Supplementary Fig. S5C) while amplification of the canonical transcript region confirmed the presence of full-length CPE mRNA across the tested conditions (Supplementary Fig. S5D). In MCF12A cells, amplicons corresponding to transcript regions predicted to encode the ∼40 kDa CPE species were relatively enriched, whereas in MCF7 spheroids, amplicons corresponding to lower-molecular-weight species were enriched instead. These findings suggest architecture- and cell-type-dependent differences in the representation of CPE transcript regions (Supplementary Fig. S5E). Amplification of a conserved CPE coding region confirmed comparable total CPE transcript levels across all conditions (Supplementary Fig. S5F). These findings suggest context-dependent differences in the representation of CPE transcript regions. However, definitive assignment of the lower-molecular-weight protein species and determination of their mechanism of generation will require transcript-level sequencing and mass-spectrometric validation.

Together, the public chromatin-occupancy data are consistent with ARNT association at the CPE locus near ERα-bound chromatin, while biochemical analyses show that ARNT loss alters the abundance and subcellular distribution of CPE-immunoreactive species in 3D spheroids. These observations provide a potential molecular context for the functional ARNT–CPE relationship and its architecture-dependent effects on spheroid growth.

### An inverse ARNT–CPE expression relationship in breast cancer patients

To examine whether the inverse expression relationship between ARNT and CPE observed in cell-line models is also detectable in patient breast tumours, we analysed publicly available transcriptomic datasets. In TCGA-BRCA Luminal A samples stratified by Hallmark Hypoxia score ^42^, ARNT and CPE expression were inversely correlated in the lowest-score tertile (n = 139, Pearson r = −0.33, P = 8.4 × 10⁻⁵; Fig. 5Aa), whereas correlations in the intermediate- and highest-score tertiles were not statistically significant (Fig. 5Ab, Ac). Within the lowest-score tertile, the inverse association remained detectable after restricting the analysis to tumours with ABSOLUTE purity > 0.60 (n = 106, Pearson r = −0.36, P = 1.8 × 10⁻⁴; Fig. 5Ad). The selective anti-correlation in low-hypoxia tumours suggests that the ARNT–CPE relationship may be most detectable under conditions of low hypoxic signalling, though whether this reflects direct masking by hypoxia-driven programmes or other confounding differences between tertiles remains untested.

To extend these observations to the single-cell level, we interrogated publicly available single-cell RNA-seq (scRNA-seq) data from primary breast tumours (GSE176078) ^43^. The proportion of CPE-expressing cancer epithelial cells was markedly lower in ER+ tumours than in TNBC and HER2+ subtypes (5.5% versus 12.1% and 46.4%, respectively; P < 0.001, Mann–Whitney U test; Fig. 5B). Within ER+ cancer epithelial cells with detectable ARNT expression (n = 1,410), ARNT-high cells exhibited significantly lower CPE expression than ARNT-low counterparts (P < 0.001, Mann– Whitney U test; Fig. 5C), consistent with the inverse ARNT–CPE relationship identified in cell-line models. Together, these findings indicate that the ARNT–CPE relationship is detectable in primary breast tumour tissue across independent datasets and at single-cell resolution, supporting its relevance beyond the experimental models described above.

**Fig. 5.**
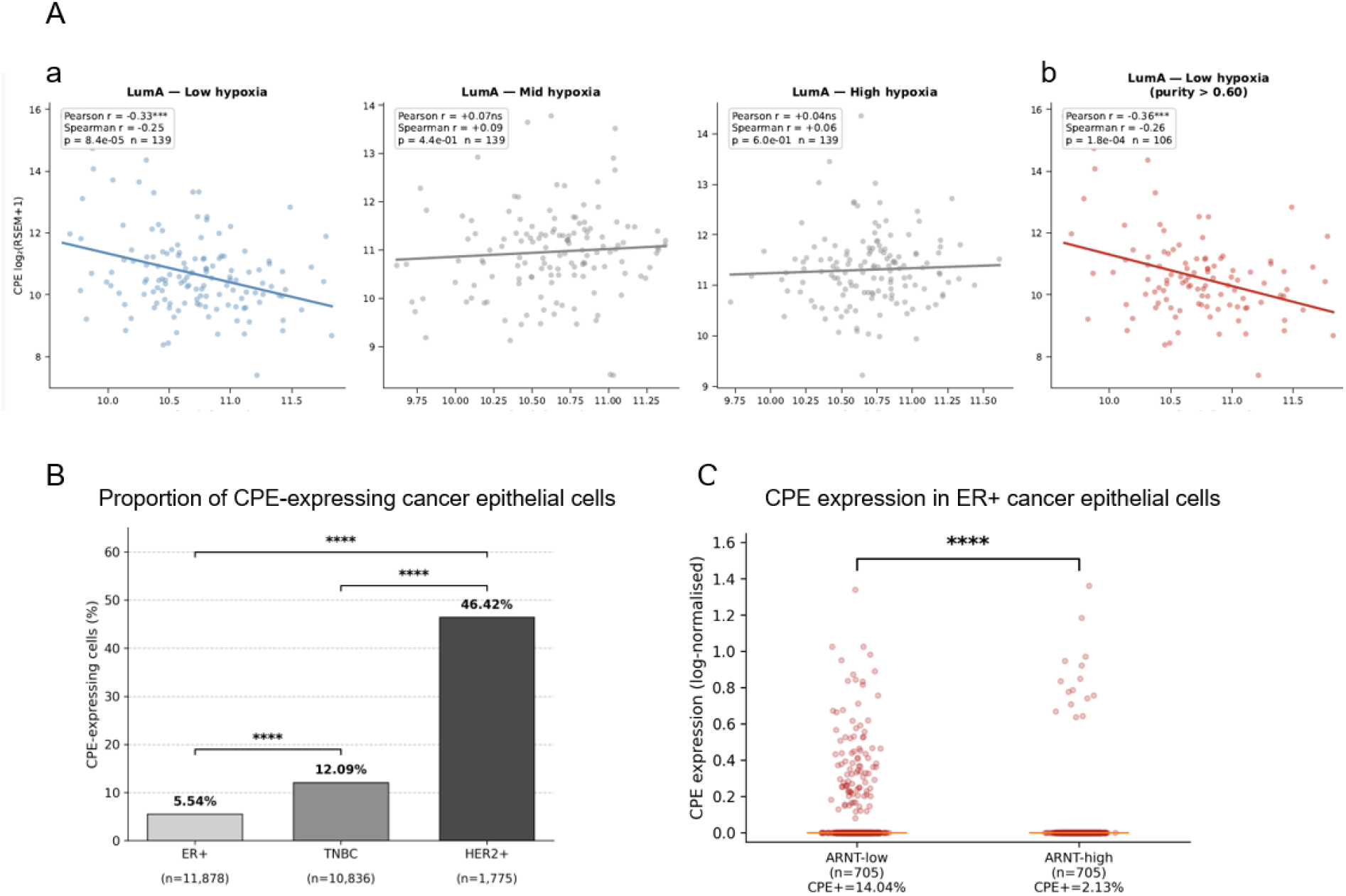
ARNT–CPE relationship in primary breast tumors. **A** An ARNT-CPE expression correlation in TCGA-BRCA Luminal A samples stratified by Hallmark Hypoxia score. Scatter plots of *ARNT* versus *CPE* expression (log₂(RSEM+1)) in TCGA-BRCA Luminal A samples stratified into low **(Aa)**, middle **(Ab)** and high **(Ac)** Hallmark Hypoxia tertiles (n = 139 per group). (Ad) Same analysis restricted to samples with ABSOLUTE tumor purity > 0.60 (n = 106). Pearson and Spearman correlation coefficients with two-sided *P* values are shown. Trend lines represent linear regression fits *\*P<0.05, **P<0.01, ***P<0.001* and *\*\*\*\*P<0.0001.* **B** Proportion of CPE-expressing cancer epithelial cells (log-normalized expression > 0) across molecular subtypes in a publicly available scRNA-seq dataset ^43^ (GSE176078; ER+, n = 11,878; TNBC, n = 10,836; HER2+, n = 1,775). Statistical comparisons by Mann–Whitney U test. \*\*\*\**P* < 0.0001. **C** CPE expression in ER+ cancer epithelial cells stratified by ARNT expression level. Cells with detectable ARNT expression (ARNT > 0, n = 1,410) were divided into ARNT-low and ARNT-high groups using the median ARNT expression value as threshold (n = 705 per group). The proportion of CPE-expressing cells (CPE+) within each group is indicated below the x-axis. Statistical comparison by Mann-Whitney U test. \*\*\*\**P* < 0.0001.

## Discussion

Our study identifies a functional ARNT–CPE axis that selectively controls 3D growth of luminal breast cancer models. Through matched 2D and 3D CRISPR screening, reciprocal ARNT perturbation, transcriptomic profiling, rescue experiments and genetic epistasis, we establish CPE as a downstream effector that is required for ARNT-loss-driven spheroid expansion and sufficient to restore growth in ARNT-overexpressing spheroids. These findings extend previous evidence that growth dependencies differ between 2D and 3D cultures ^11,12^ by showing that ARNT loss has opposing effects on monolayer and spheroid growth and provide evidence that architecture can invert, rather than simply alter the magnitude of a transcription factor-dependent fitness phenotype.

The most striking feature of our data is not merely that ARNT regulates different transcriptional programmes in 2D and 3D, but that its net effect on cellular fitness is directionally inverted between these conditions. ARNT is best known as the obligate dimerization partner for HIF-1α and AhR ^17–19^, and its activity is strongly influenced by partner availability and cellular signalling context ^20,21^. Our findings extend this view in an unexpected direction: *ARNT* loss reduces growth in monolayer culture but promotes spheroid expansion, a phenotype reproduced across two luminal breast cancer cell lines and functionally linked to CPE in 3D architecture.

In monolayer culture, *ARNT* loss was accompanied by reduced expression of survival- and growth-associated genes, including TERT, SLC7A5 and NFE2L2, and induction of stress-response and cell-cycle arrest genes, including CDKN1A, BTG2, NUPR1 and DDIT3. By contrast, the transcriptional response to ARNT perturbation was more restricted in spheroids, with approximately six-to eightfold fewer differentially expressed genes and reciprocal regulation of CPE among the most prominent 3D-selective changes. This pattern indicates that the same transcription factor can perform distinct regulatory roles depending on tissue architecture — a property suggested for chromatin-context-dependent regulators ^2,4^ but that has not, to our knowledge, been resolved at this resolution for a single transcription factor in matched 2D/3D systems in breast cancer. The growth-suppressive effect of ARNT overexpression in 3D may additionally reflect induction of *TGFB2* (log_2_FC = 1.91, adjusted *P* = 0.042), which showed the largest fold change among upregulated genes other than *ARNT* itself, together with increased AHRR expression which was modestly but significantly induced (log_2_FC = 0.80, adjusted *P* = 0.035) (Supplementary Table S9D). These changes identify additional ARNT-responsive genes, but their contribution to spheroid growth remains to be established.

Several aspects of CPE biology are compatible with an architecture-dependent role in spheroid growth. CPE has established roles in neuropeptide processing and trophic factor secretion, and secreted CPE has been reported to activate pro-survival signalling in tumour contexts ^25^. A nuclear-localised N-terminal truncated form of CPE (CPE-ΔN) ^26^ has also been reported in human cancer cells and implicated in transcriptional regulation and metastasis-related gene expression. Whether nuclear CPE activity contributes to the architecture-dependent phenotypes observed in breast cancer spheroids remains to be determined. The selective engagement of CPE as an effector in 3D may reflect the altered secretory demands of spheroid organisation, where paracrine signalling and vesicle trafficking operate under different spatial constraints than in monolayer.

Publicly available ChIP–seq data provided a possible regulatory context for the ARNT– CPE relationship. ARNT ChIP–seq peaks were detected across the CPE locus in proximity to ESR1-occupied chromatin, while the absence of a canonical E-box motif and the presence of a half-ERE near the transcription start site were consistent with the possibility of ERα-associated recruitment rather than canonical ARNT DNA recognition. These observations do not establish direct ARNT binding or ERα-mediated tethering in our experimental models, but they identify a plausible chromatin context through which ARNT could influence CPE expression. Primary tumour analyses further supported the context dependence of this relationship. ARNT and CPE were inversely correlated in the lowest Hallmark Hypoxia score tertile of TCGA-BRCA Luminal A tumours, while single-cell analysis identified lower CPE expression in ARNT-high ER+ cancer epithelial cells. The markedly lower proportion of CPE-expressing epithelial cells in ER+ tumours compared with TNBC and HER2+ tumours further suggests that CPE regulation differs across breast cancer subtypes. Together, these observations support the relevance of the inverse ARNT–CPE relationship in primary ER+ disease without implying that the same regulatory state is maintained across all tumour contexts.

Alongside CPE, transcriptomic analysis identified LXN as a second ARNT-responsive candidate with an opposing effect on spheroid growth. LXN loss increased spheroid area, whereas CPE loss reduced it. LXN is an endogenous carboxypeptidase inhibitor with reported tumour-suppressive functions ^47^; however, its deleterious mutation and lack of detectable expression in T47D cells prevented its evaluation as a shared effector across both luminal models. Future studies in LXN-intact systems may determine whether CPE and LXN participate in parallel or opposing protease-regulatory programmes downstream of ARNT.

Our screening strategy focused on chromosome 1q, one of the most frequently gained chromosome arms in breast cancer. Broad or partial 1q gain was detected in 92% of the breast cancer cell lines examined in CCLE ^44^. The use of MCF7 and T47D cells provided complementary 1q copy-number contexts, while the absence of a spheroid phenotype following targeting of a nearby intergenic region supports the conclusion that the observed effects arise from ARNT gene perturbation rather than nonspecific regional effects associated with chromosome 1q targeting. Moreover, the promotion of spheroid growth by *ARNT* loss was also observed in MCF12A cells, which do not share the same transformed luminal background. Thus, the architecture-dependent component of ARNT function is not solely explained by the genomic configuration of chromosome 1q, although its relevance across broader breast cancer genotypes remains to be determined.

Several points should be considered when interpreting these findings. The methylcellulose-based spheroid system used here captures key architectural features, including three-dimensional compaction and diffusion gradients, but does not reproduce the full tumour microenvironment, including stromal, vascular and immune components. In addition, the molecular identity of the nuclear CPE-immunoreactive species detected in 3D spheroids remains unresolved and will require mass-spectrometric validation. Finally, functional validation was performed in two luminal breast cancer cell lines and one non-tumorigenic model; systematic testing in broader luminal breast cancer panels, patient-derived organoids and *in vivo* models will be important to define the generality and disease relevance of the ARNT–CPE axis.

ARNT and its growth-restrictive effect in 3D architecture was observed in both luminal breast cancer cells and the non-tumorigenic MCF12A model. From a translational perspective, the requirement for ARNT in 2D growth was observed in MCF7 and T47D cells but not in MCF12A cells, suggesting that the therapeutic consequences of targeting ARNT would be strongly context dependent. CPE may represent a more experimentally tractable component of this axis because of its enzymatic and secretory functions, although its multiple intracellular, extracellular and lower-molecular-weight forms complicate direct therapeutic targeting. Its reported roles in tumour cell survival ^25^, together with the ability of CPE loss to suppress ARNT-loss-driven spheroid expansion, support further investigation of the ARNT–CPE axis in patient-derived models.

Architecture-dependent inversion of transcription factor function carries implications beyond ARNT. Our data argue that for transcription factors with multiple partner choices, a 3D context can not only modulate the magnitude of a fitness phenotype but invert its direction. In a 2D-only screen, ARNT would have been classified as broadly pro-tumorigenic, and CPE would not have been detected as a selective effector at all— readings inconsistent with the 3D phenotype. This supports the systematic use of paired 2D/3D designs for the discovery of architecture-dependent vulnerabilities, particularly for context-sensitive regulators such as bHLH–PAS proteins, steroid receptors or YAP/TAZ-class effectors. In summary, our study identifies ARNT as an architecture-dependent regulator of luminal breast cancer growth, whose fitness function is inverted between monolayer and spheroid conditions and CPE acts as a functional mediator of ARNT-loss-associated spheroid expansion. By resolving this inversion in matched 2D and 3D systems, we provide a framework for identifying context-sensitive dependencies that are misassigned in conventional monolayer screens.

## Supplementary Information

**Supplementary File 1 (.xlsx):** Supplementary tables. Library annotations, primer sequences, copy-number summaries, CRISPR screening results and RNA-seq differential expression results.

**Supplementary File 2 (.pdf):** Supplementary figures and figure legends. Supporting analyses and experimental validation accompanying the main figures.

## Declarations

### Ethics approval and consent to participate

This study was conducted exclusively using established human cell lines obtained from ATCC. No human subjects, human tissue, or animal experiments were involved. Ethical approval was therefore not applicable.

### Consent for publication

This study was not contained data from any individual person. Consent for publication was therefore not applicable.

### Availability of data and materials

Raw and processed RNA-seq data and CRISPR screen sgRNA count matrices and analysis outputs have been deposited at the Gene Expression Omnibus (GEO) as a superseries under accession number GSE333963. CCLE ABSOLUTE copy-number data are publicly available at https://depmap.org. ChIP–seq peak collections used in this study are publicly available at https://chip-atlas.org. TCGA-BRCA RNA-seq and clinical data are available at https://portal.gdc.cancer.gov. All other data supporting the findings of this study are available within the article and its Supplementary Information files, or from the corresponding author upon reasonable request.

### Authors’ Contributions

GK and HOC designed and conceptualized all experiments. HOC was responsible for all functional experiments including CRISPR screening and validation assays. FBB was responsible for bioinformatic analyses including CRISPR screen, RNA-seq, and ChIP-seq, and performed the generation of bioinformatic figures. HOC and EK established and performed CRISPR screens and conceptualized validation experiments. ENÇ, MSO, and EG performed functional experiments. ZC performed western blots. BD analyzed scRNA-seq data and generated related figures. DT and GK supervised this study and provided critical intellectual input throughout. GK acquired funding. HOC and GK wrote and edited the manuscript.

## Acknowledgements

The authors gratefully acknowledge use of the services and facilities of the Koç University Research Center for Translational Medicine (KUTTAM), funded by the Presidency of Turkey, Presidency of Strategy and Budget. The content is solely the responsibility of the authors and does not necessarily represent the official views of the Presidency of Strategy and Budget. The authors thank Eda Yıldırım (Koç University) for critical reading of the manuscript and valuable feedback. RNA sequencing services were provided by Genera Biotech (Refgen, Ankara, Turkey).

