## Supplementary Figures for "Tumour architecture inverts ARNT-dependent growth regulation and reveals CPE-mediated spheroid expansion in luminal breast cancer"

Hande Ozkan-Coskun *et al.*

\*Corresponding author.

.

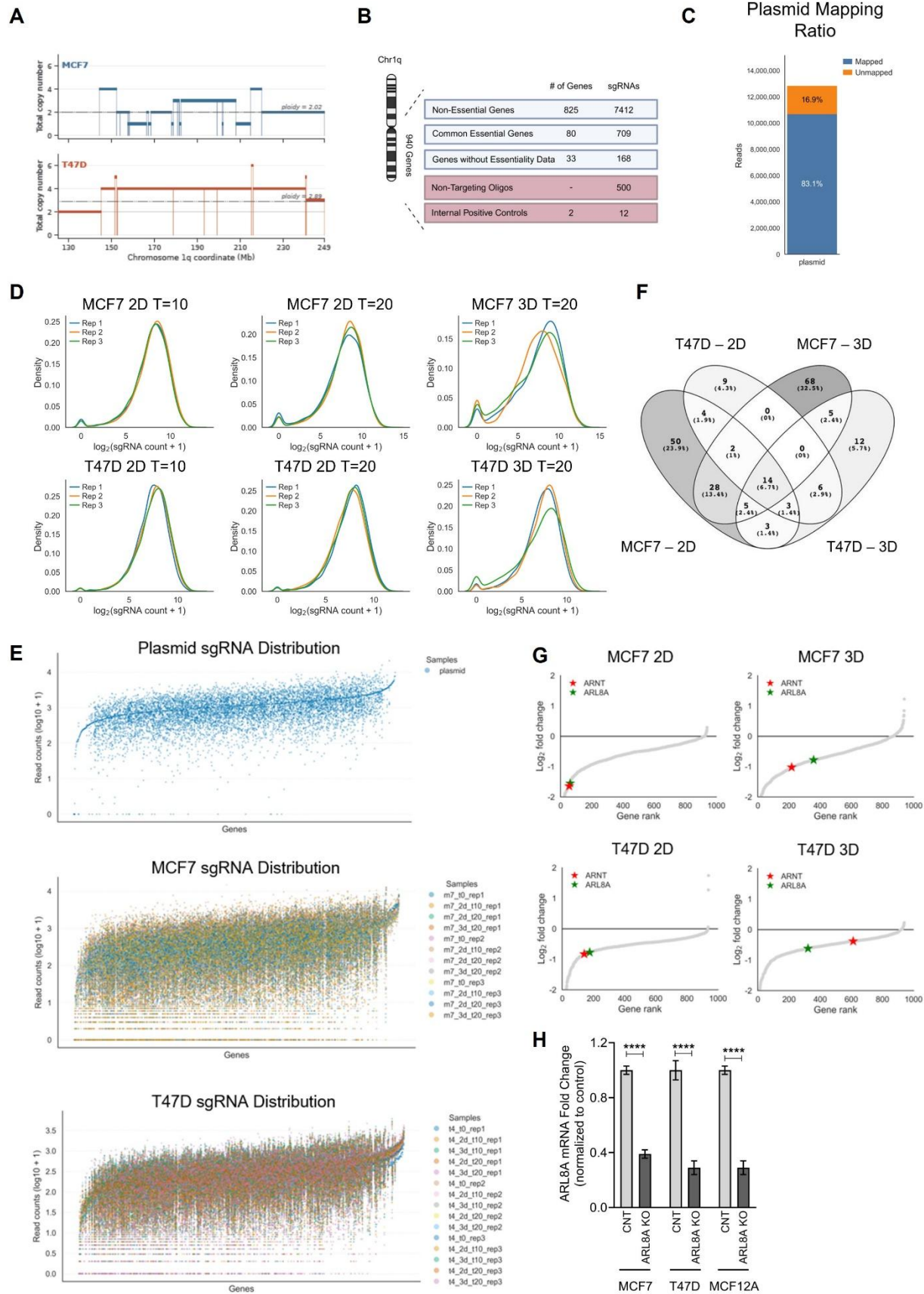

**Supplementary Figure S1. Design, quality control, and candidate selection of the pooled CRISPR screen.**

**(A)** Chromosome 1q absolute copy number profiles of MCF7 and T47D cells from CCLE ABSOLUTE data [Carter et al., 2012]. Each horizontal segment represents a contiguous region with constant inferred total copy number. Sample ploidy is indicated by the dotted line (MCF7 = 2.02, T47D = 2.89); the diploid reference level (CN = 2) is shown in grey. MCF7 carries focal Chr1q gains on a near-diploid background, whereas T47D shows broad Chr1q amplification on a genome-doubled background, providing two complementary genomic contexts. **(B)** Schematic representation of the LIBGENE1Q library composition. The library targets Chr1q-encoded genes with 8,250 sgRNAs against 825 non-essential genes, 800 sgRNAs against 80 common essential genes, 200 sgRNAs against 2 internal positive control genes, and 500 non-targeting (NT) sgRNAs. **(C)** Quality control of the plasmid library showing the proportion of mapped and unmapped reads (83.1% mapped). The high mapping rate confirms successful library construction and preserved sgRNA representation prior to lentiviral packaging. **(D)** Density distribution plots of log<sub>2</sub>-transformed sgRNA counts across biological replicates and time points for MCF7 (2D T=10, 2D T=20, 3D T=20) and T47D (2D T=10, 2D T=20, 3D T=20) cells. Overlapping distributions across replicates confirm inter-replicate reproducibility at each experimental time point. **(E)** Whole-library sgRNA representation plots showing log<sub>10</sub>-transformed read counts per gene across all samples in the plasmid library (top), MCF7 (middle) and T47D (bottom) screens. Each point represents an individual sgRNA colored by sample identity. The uniform distribution across gene ranks confirms broad sgRNA representation and minimal dropout prior to and during screening. **(F)** Venn diagram showing the overlap of significantly depleted candidate genes identified across MCF7 2D, MCF7 3D, T47D 2D and T47D 3D dropout screens (FDR < 0.1, log<sub>2</sub>FC < -0.8). **(G)** Rank-based gene prioritization plots showing log<sub>2</sub> fold-change distributions for the 3D versus 2D comparison in MCF7 2D, MCF7 3D, T47D 2D and T47D 3D conditions. ARNT (red star) and ARL8A (green star) are highlighted as candidate architecture-dependent genes showing relative enrichment in 3D compared with 2D. **(H)** RT-qPCR validation of ARL8A knockout efficiency in MCF7, T47D and MCF12A cells under 2D monolayer culture conditions. ARL8A mRNA levels were normalized to Beta-actin and presented relative to the corresponding control group.

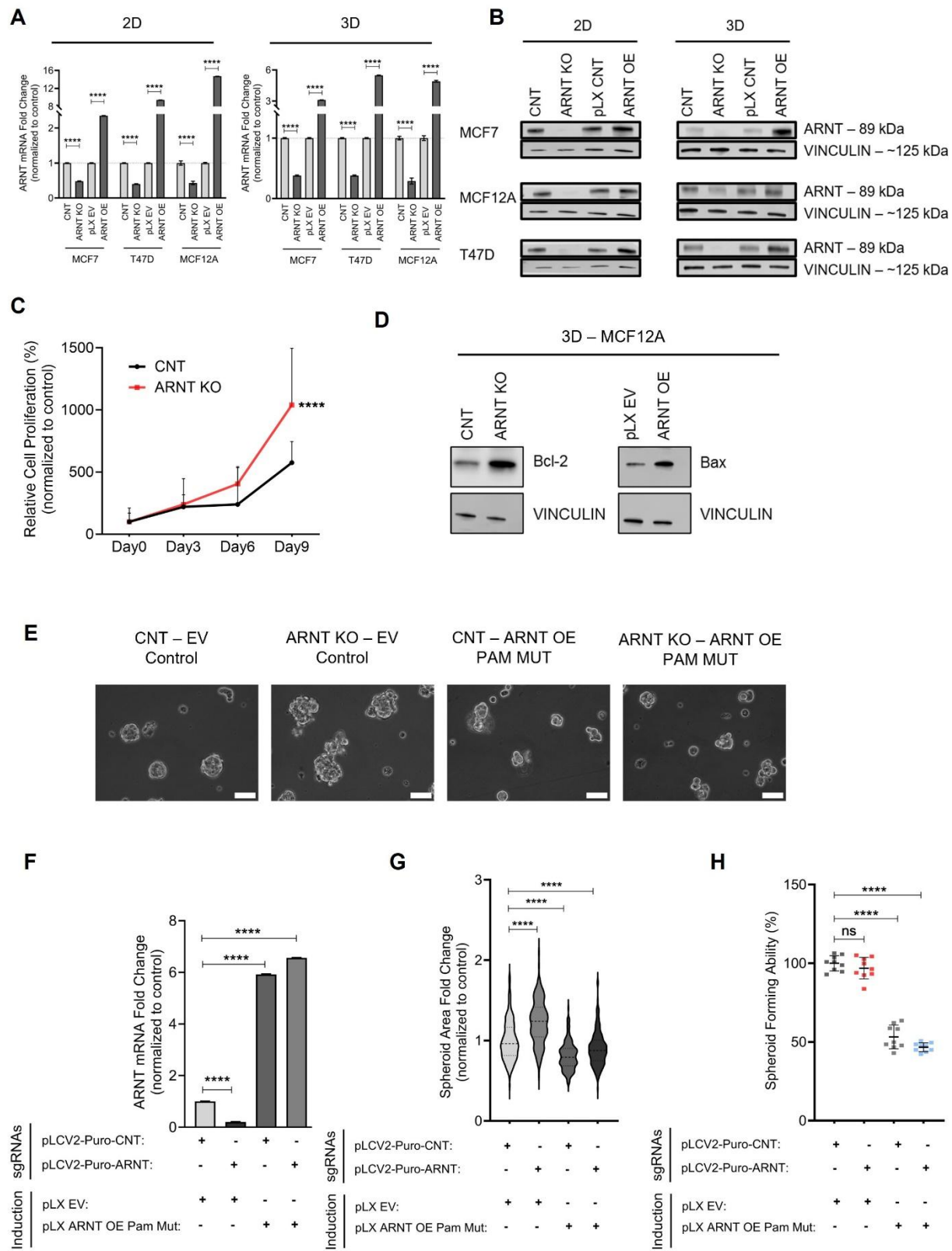

**Supplementary Figure S2. Validation of ARNT perturbation and rescue experiments.**

**(A)** RT-qPCR validation of ARNT knockout (KO) and overexpression (OE) efficiency in MCF7, T47D and MCF12A cells under 2D monolayer and 3D spheroid culture conditions. ARNT mRNA levels were normalized to Beta-actin and presented relative to the corresponding control group. **(B)** Immunoblot validation of ARNT protein expression following *ARNT* KO and OE in MCF7, T47D and MCF12A cells under 2D and 3D conditions; vinculin, loading control. **(C)** CellTiter-Glo 3D viability assay in MCF7 spheroid cultures showing increased viable cell mass following *ARNT* KO compared with control over the indicated time points. **(D)** Immunoblot analysis of Bcl-2 and Bax protein levels in MCF12A 3D spheroids following ARNT KO and ARNT OE; vinculin, loading control. **(E)** Representative brightfield images of MCF7 spheroids under control (CNT–EV), *ARNT* KO (ARNT KO–EV), CNT–ARNT OE PAM-mutant and ARNT KO–ARNT OE PAM-mutant rescue conditions; scale bar, 70 μm. **(F)** RT-qPCR validation of *ARNT* expression in rescue experiments confirming efficient induction of the PAM-mutant ARNT overexpression construct. ARNT mRNA levels were normalized to Beta-actin and presented relative to the corresponding control group. **(G)** Quantification of spheroid area fold change across CNT, *ARNT* KO and ARNT rescued conditions. Re-expression of PAM-mutant ARNT restored the spheroid growth phenotype induced by *ARNT* loss. **(H)** Quantification of spheroid-forming ability across the indicated groups. Data represent mean ± s.e.m. from three biological replicates. $P < 0.05$ (\*), $P < 0.01$ (\*\*), $P < 0.001$ (\*\*\*) and $P < 0.0001$ (\*\*\*\*); ns, not significant; two-tailed Student's *t*-test.

**A**

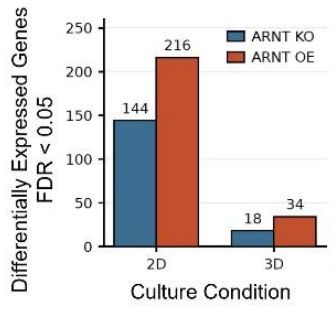

**B**

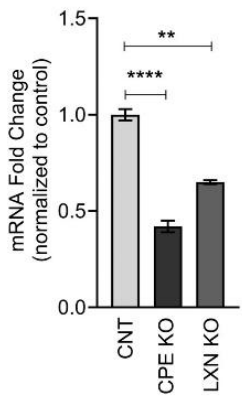

**C**

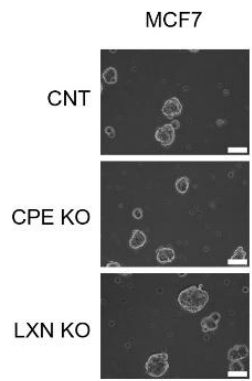

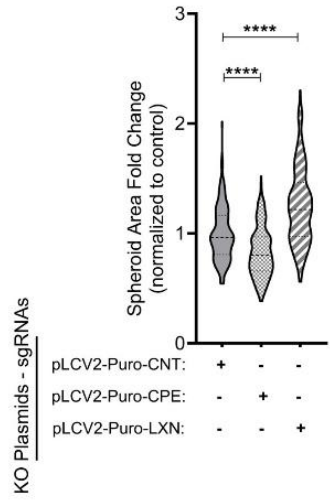

**D**

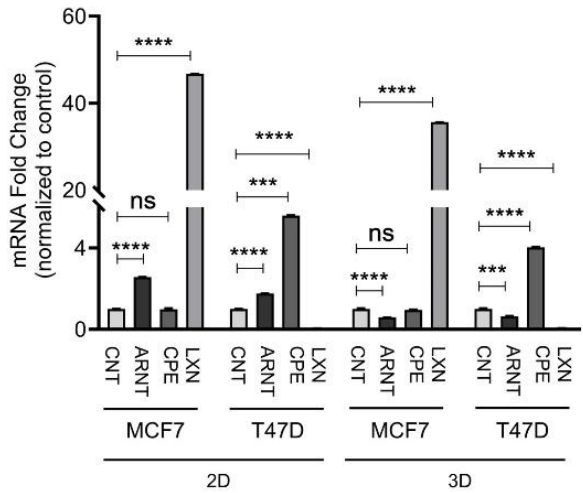

**Supplementary Figure S3. Validation and functional analysis of the ARNT–CPE/LXN axis in 2D and 3D breast cancer models.**

**(A)** Number of differentially expressed genes (DEGs; FDR < 0.05) identified by DESeq2 analysis of *ARNT* KO and *ARNT* OE conditions relative to non-targeting control under 2D and 3D culture in MCF7 cells. The 6–8-fold reduction in DEG counts in 3D relative to 2D indicates substantial attenuation of the transcriptional response to *ARNT* perturbation by spheroid architecture. **(B)** Confirmation of *CPE* and *LXN* knockout efficiency after targeting each gene separately by RT-qPCR in MCF7 3D spheroid cultures. mRNA levels were normalized to Beta-actin and presented relative to the corresponding control group. **(C)** Representative brightfield images of MCF7 3D spheroids following *CPE* KO and *LXN* KO (left); scale bar, 70 μm. Quantification of spheroid area fold change (right): *CPE* depletion reduced spheroid growth, whereas *LXN* knockout increased spheroid area. **(D)** RT-qPCR analysis of *ARNT*, *CPE* and *LXN* expression in MCF7 and T47D cells cultured in 2D and 3D, normalized to MCF12A. *ARNT* expression decreased in 3D spheroids while *CPE* showed a modest increase. *LXN* was undetectable in T47D. mRNA levels were normalized to Beta-actin and presented relative to the corresponding control group. Data represent mean ± s.e.m. from three biological replicates. $P < 0.05$ (\*), $P < 0.01$ (\*\*), $P < 0.001$ (\*\*\*) and $P < 0.0001$ (\*\*\*\*); ns, not significant; two-tailed Student's *t*-test.

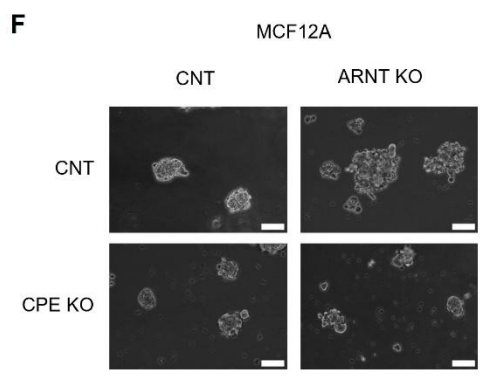

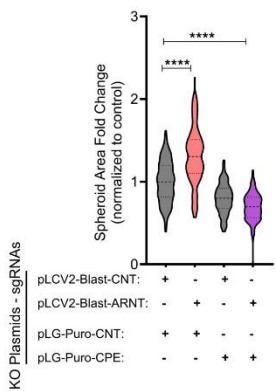

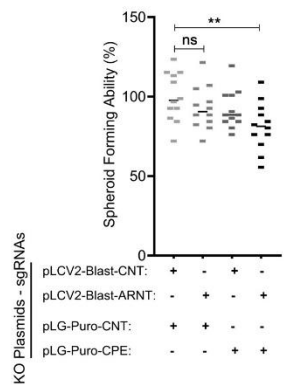

**Supplementary Figure S4. Conservation of the ARNT–CPE epistatic relationship in T47D and MCF12A spheroids.**

**(A)** RT-qPCR validation of combinatorial loss-of-function in MCF7 3D spheroids confirming efficient single and dual perturbation of *ARNT* and *CPE*. **(B)** RT-qPCR validation of combinatorial overexpression in MCF7 3D spheroids confirming successful induction of *ARNT* and *CPE* constructs in single and combined conditions. RT-qPCR validation of combinatorial *ARNT* and *CPE* perturbations in **(C)** T47D and **(D)** MCF12A confirming efficient depletion of *ARNT*, *CPE* and *ARNT–CPE* double-knockout conditions following CRISPR-based targeting. mRNA levels were normalized to Beta-actin and presented relative to the corresponding control group. **(E)** Representative brightfield images of T47D 3D spheroids under combinatorial *ARNT* and *CPE* perturbation (left); scale bar, 70 μm. Quantification of spheroid area fold change (center) and spheroid-forming ability (right). *ARNT* KO increased spheroid size, whereas concurrent *CPE* targeting reduced spheroid growth and spheroid-forming ability. **(F)** Representative brightfield images of MCF12A 3D spheroids following *ARNT* and *CPE* co-perturbation (left); scale bar, 70 μm. Quantification of spheroid area fold change (center) and spheroid-forming ability (right). Data represent mean ± s.e.m. from three biological replicates. $P < 0.05$ (\*), $P < 0.01$ (\*\*), $P < 0.001$ (\*\*\*) and $P < 0.0001$ (\*\*\*\*); ns, not significant; two-tailed Student's *t*-test.

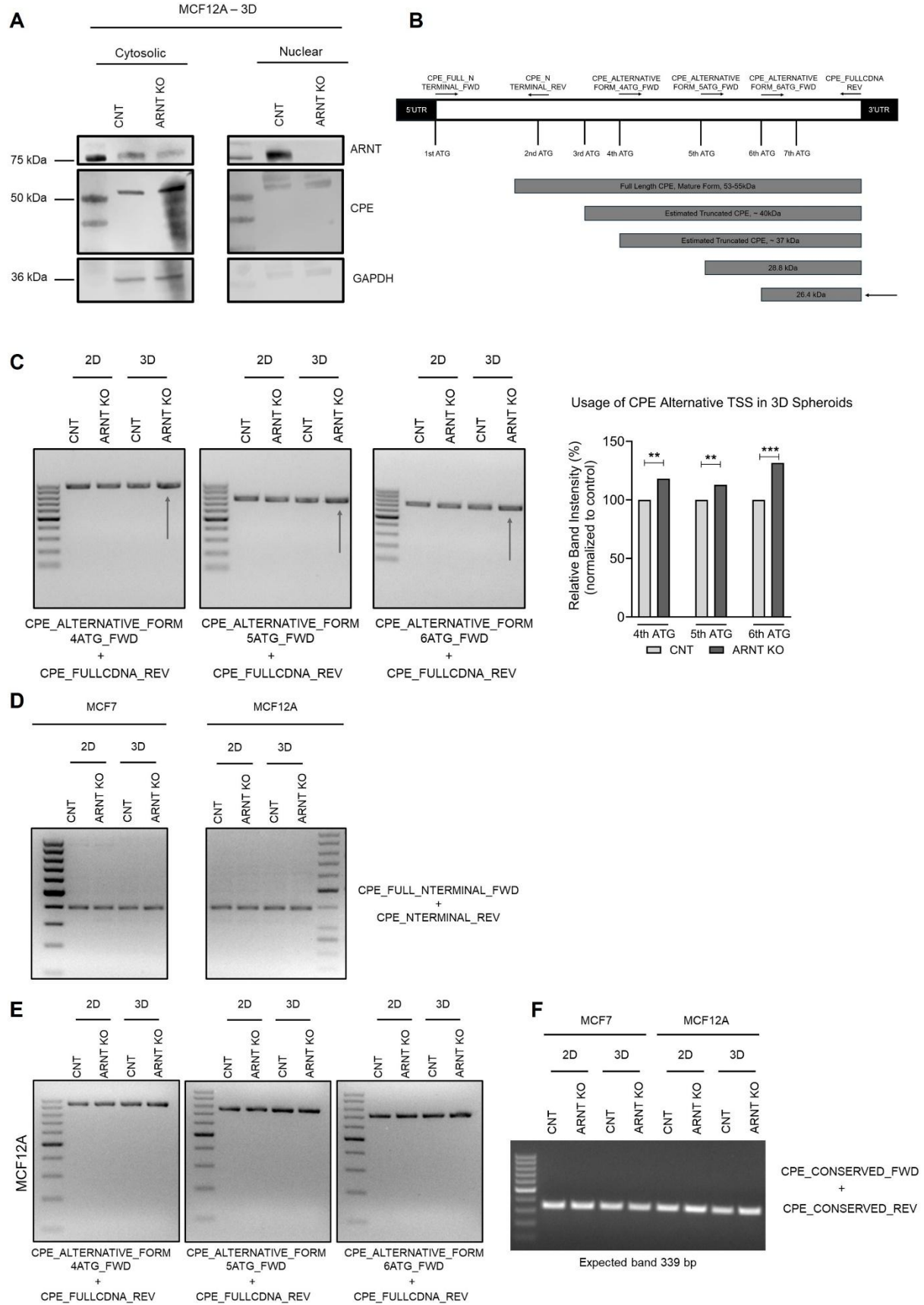

**Supplementary Figure S5. Western blot and PCR-based analysis of CPE protein and transcript architecture.**

**(A)** Subcellular fractionation followed by western blot analysis in MCF12A-derived 3D spheroids. Lower molecular weight CPE species (~27 kDa) were not detected in non-tumorigenic MCF12A ARNT knockout cells under 3D growth conditions. These observations suggest that the emergence of this species is dependent on both tumor context and three-dimensional growth conditions. Molecular weight markers are shown in the leftmost lane of each gel. **(B)** Schematic representation of the *CPE* transcript structure highlighting potential alternative translation initiation sites and the predicted protein products arising from differential ATG usage. Canonical full-length CPE (53–55 kDa) and alternative forms (~40 kDa, ~37 kDa, ~28.8 kDa, and ~26.4 kDa) are indicated. **(C)** PCR amplification of CPE coding regions downstream of alternative translation initiation sites in MCF7 cells under 2D and 3D culture conditions with control (CNT) and ARNT knockout (ARNT KO) groups. Primer pairs were designed to initiate amplification at the 4th, 5th, and 6th ATG codons of the CPE transcript, respectively, each paired with a common reverse primer (CPE\_FULLCDNA\_REV). Increased amplification from the 4th, 5th and 6th ATG-initiated regions was observed selectively in ARNT KO 3D spheroids (arrows), consistent with preferential usage of downstream translation initiation sites under these conditions and concordant with the enrichment of lower molecular weight CPE species observed by immunoblot. **Left:** CPE\_ALTERNATIVE\_FORM\_4ATG\_FWD + CPE\_FULLCDNA\_REV, expected amplicon size 952 bp. **Middle:** CPE\_ALTERNATIVE\_FORM\_5ATG\_FWD + CPE\_FULLCDNA\_REV, expected amplicon size 760 bp. **Right:** CPE\_ALTERNATIVE\_FORM\_6ATG\_FWD + CPE\_FULLCDNA\_REV, expected amplicon size 700 bp. The 100 bp DNA ladder is shown in the leftmost lane of each gel. Data represent mean ± s.e.m. from three biological replicates. $P < 0.05$ (\*), $P < 0.01$ (\*\*), $P < 0.001$ (\*\*\*) and $P < 0.0001$ (\*\*\*\*); ns, not significant; two-tailed Student's *t*-test. **(D)** PCR amplification using a primer pair spanning the canonical translation initiation region of *CPE* (CPE\_FULL\_NTERMINAL\_FWD + CPE\_NTERMINAL\_REV; expected amplicon size: 388 bp). A single amplification product corresponding to the full-length *CPE* transcript was consistently detected across all experimental conditions in both MCF7 and MCF12A cells under 2D and 3D cultures. No substantial differences in PCR product intensity were observed between MCF7 cells and non-tumorigenic MCF12A cells, indicating that the canonical full-length *CPE* transcript remains broadly conserved consistent with the blotting results. The 100 bp DNA ladder indicates the PCR product was around 400bp. **(E)** Representative agarose gel images showing PCR amplification patterns obtained from MCF12A cells under the indicated experimental conditions. Distinct intensity after amplification was observed across primer sets, indicating the possibility of differential usage of internal translation initiation regions. The enhanced abundance of 952 bp transcript in MCF12A-derived 3D spheroids upon ARNT loss, compared with the shorter transcript variants predominantly observed in MCF7 cells, was consistent with the western blot findings demonstrating marked enrichment of the ~40 kDa and/or ~37 kDa CPE protein species following ARNT depletion in MCF12A 3D models. 100 bp DNA ladder indicates the PCR products were around 1kb and around 800 bp. **(F)** Amplification of a conserved *CPE* coding region using CPE\_CONSERVED\_FWD and CPE\_CONSERVED\_REV (primer pair #5) demonstrating stable transcript detection, serving as a reference for total *CPE* transcript abundance across all experimental groups with expected product length 339 bp.

### **Supplementary Tables**

**Supplementary Table S1. sgRNA sequences in the LIBGENE1Q library.** Full list of sgRNA sequences included in the LIBGENE1Q CRISPR library used for parallel 2D and 3D screens.

**Supplementary Table S2. Primer pairs used for library amplification by two-step PCR.** Sequences of primers used in the two-step PCR protocol for sgRNA library preparation and amplification.

**Supplementary Table S3. Oligonucleotide sequences used in functional experiments.** Sequences of all oligonucleotides, used in functional perturbation experiments.

**Supplementary Table S4. Primer sequences used in cloning experiments.** Sequences of primers used for cloning of CPE overexpression and PAM mutated knockout plasmid constructs.

**Supplementary Table S5. Primer sequences used for RT-qPCR.** Sequences of forward and reverse primers used for RT-qPCR quantification of gene expression, including target genes and housekeeping controls.

**Supplementary Table S6. Primer pairs used for PCR-based detection of CPE transcript variants.** Sequences of forward and reverse primers used to amplify CPE transcript regions.

**Supplementary Table S7. ABSOLUTE characterization of breast cancer cell lines from CCLE.** Per-cell-line summary of absolute copy number features for 52 breast cancer cell lines profiled by the ABSOLUTE algorithm [REF] in CCLE (release 2018-12-27). For each cell line, the table lists ABSOLUTE-derived purity, ploidy and number of inferred genome doublings, together with Chr1q-specific metrics: number of segments overlapping Chr1q (defined as Chr1 coordinates $\geq 125$ Mb on hg19), median and length-weighted mean total copy number, maximum and minimum total copy number, percentage of Chr1q with copy number above ploidy (gain) or below ploidy (loss), and overall Chr1q status classification (broad gain, $\geq 50\%$ gained; partial gain, 20–50%; broad loss, $\geq 50\%$ lost; partial loss, 20–50%; no major change otherwise). MCF7 and T47D, the cell lines used in this study, are highlighted.

**Supplementary Table S8. CRISPR screen results.** sgRNA count matrices and screen analysis outputs for 2D and 3D conditions, including gene-level fitness scores and ranking.

**Supplementary Table S9. Differentially expressed genes from RNA-seq analysis.** Gene-level differential expression values for all pairwise contrasts (KO vs. NT and OE vs. NT) under 2D and 3D conditions in MCF7 cells, including log2 fold change, adjusted p-value and expression statistics.
